# Golden hamster piRNAs are necessary for early embryonic development and establishment of spermatogonia

**DOI:** 10.1101/2021.01.27.428513

**Authors:** Zuzana Loubalova, Helena Fulka, Filip Horvat, Josef Pasulka, Radek Malik, Michiko Hirose, Atsuo Ogura, Petr Svoboda

## Abstract

PIWI-associated RNAs (piRNAs) support the germline by suppressing retrotransposons and genes. In mice, piRNAs are essential for spermatogenesis but not oogenesis. To test how this applies to other mammals, we deleted *Mov10l1* helicase in golden hamster, whose piRNA pathway is configured more similarly to that of other mammals. *Mov10l1^−/−^* male hamsters showed impaired establishment of spermatogonia accompanied by transcriptome dysregulation and a surge in MYSERV retrotransposon expression. The rare viable spermatogenic cells showed a meiotic failure phenotype like *Mov10l1^−/−^* mice. Female *Mov10l1^−/−^* hamsters were sterile due to post-meiotic loss of developmental competence in zygotes. Unique phenotypes of *Mov10l1^−/−^* hamsters demonstrate the adaptive nature of piRNA-mediated control of genes and retrotransposons in order to confront emerging genomic threats or acquire new physiological roles.

## INTRODUCTION

The PIWI-associated RNA (piRNA) pathway is a key germline-specific silencing mechanism crucial for defending genome integrity against transposable elements (TEs) [reviewed in (1,2)]. Mammalian primary piRNAs originate from specific loci (piRNA clusters) as long precursor transcripts, which interact with the essential and conserved helicase MOV10L1, which feeds precursor transcripts into piRNA biogenesis mechanism (3–5). Mammalian piRNAs fall into four categories: 1) 26-28 nt retrotransposon-derived piRNAs produced mainly in fetal testes, 2) 26-27 nt postnatal piRNAs from non-repetitive sequences including 3’ ends of mRNAs, 3) 29-30 nt mostly non-repetitive highly abundant pachytene piRNAs produced from ~100 loci in spermatocytes and spermatids, and 4) oocyte-specific ~20 nt piRNAs enriched in antisense sequences of recently evolved TEs (6–10). Mature piRNAs associate with the PIWI subfamily of Argonaute proteins (11). Mouse (*Mus musculus*), the leading mammalian model for the piRNA pathway, utilizes three PIWI proteins: PIWIL1 (MIWI), PIWIL2 (MILI), and PIWIL4 (MIWI2). Loss of MOV10L1 or PIWI proteins revealed specific essential roles of piRNAs in spermatogenesis but not in oogenesis (3,4,12–14). The dispensability of piRNAs in females could be a mouse-specific feature. While piRNAs inhibit retrotransposon expression during mouse oogenesis (15,16), the effective RNA interference (RNAi) pathway, which specifically evolved in mice (17), functionally overlaps with the maternal piRNA pathway (18). Furthermore, mice lack PIWIL3, which binds ~20 nt piRNAs and is expressed in bovine and human oocytes (10,19).

Understanding conserved and derived aspects of the mammalian piRNA pathway requires proper comparative analysis. As an optimal comparative model we selected the golden hamster *Mesocricetus auratus*. Despite ~24 million years of independent evolution (20), the hamster shares many anatomical and physiological features with the mouse, including fast zygotic genome activation, short gestation, and large litter size (21). Importantly, golden hamster retained four PIWI paralogs (Fig. 1A), its oocytes likely lack highly active RNAi (22) and it is amenable to genetic manipulations (23).

**Figure 1.**
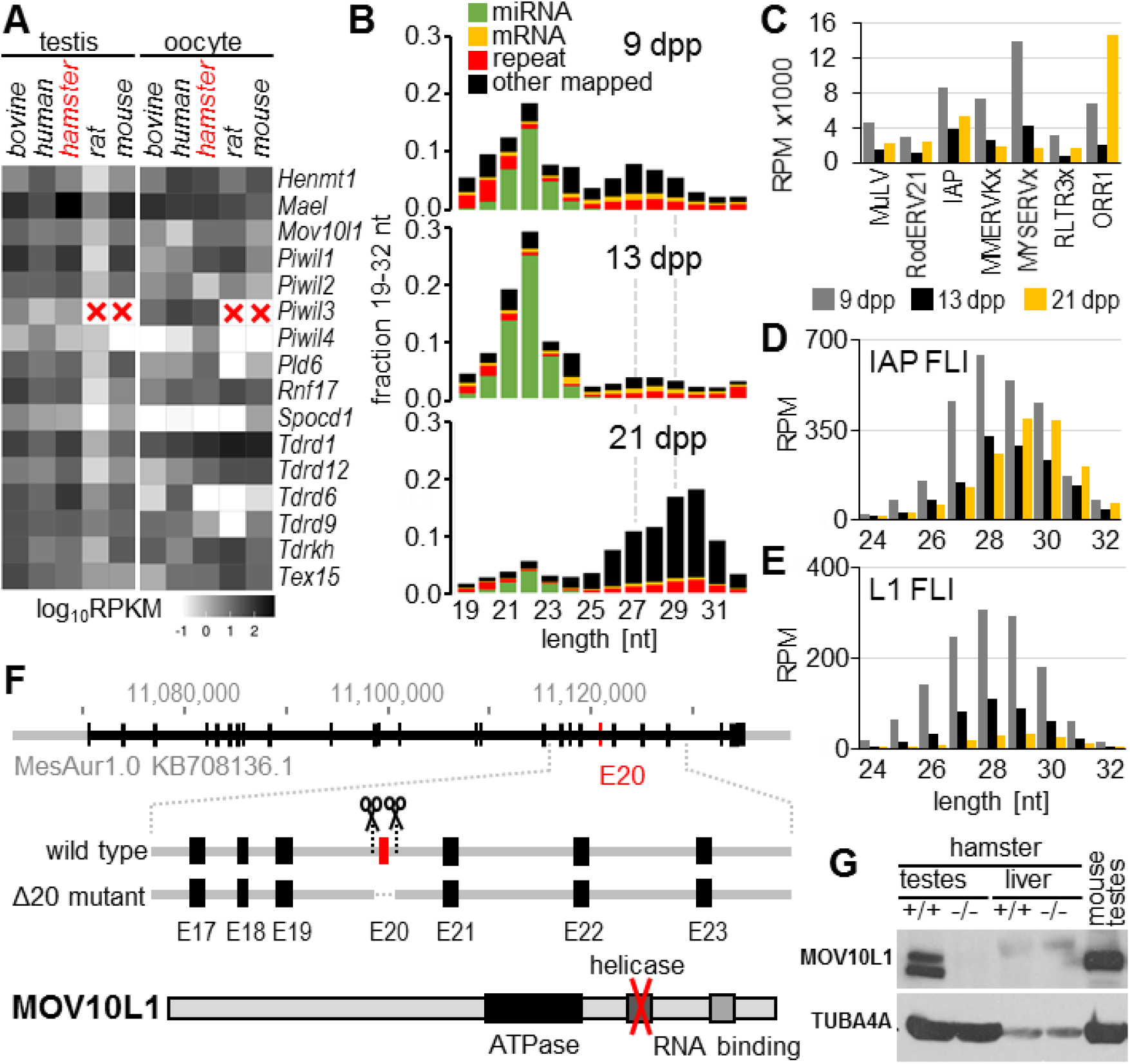
Golden hamster piRNA pathway and the *Mov10l1* knock-out. (A) Expression of piRNA pathway factors in testes and oocytes of five mammals. Mouse and rat lack *Piwil3*. (B) Distribution of 19-32 nt long RNAs from testes at 9, 13, and 21 dpp. Average of two biological replicates. (C) 24-32 nt small RNAs mapping to LTR retrotransposon groups selected for low nucleotide exchange rate and high 24-31 nt RNA abundance (see Supplementary text for details). The y-axis displays reads per million (RPM) of 19-32 nt sequences. (D, E) Distribution of 24-32 nt small antisense RNAs from testes at 9, 13, and 21 dpp that perfectly mapped to FLI IAP or L1 insertions, respectively. The y-axis displays RPM of 19-32 nt sequence reads. (F) MOV10L1 protein organization and knock-out strategy. Scissors depict CRISPR/Cas9 cleavage positions flanking exon 20 (red rectangle). (G) Western blotting shows loss of MOV10L1 in mutant testes. Other samples were a negative control (liver) and positive control (mouse testes) for antibody specificity. Two bands in hamster testes imply two MOV10L1 isoforms.

## RESULTS & DISCUSSION

### Characterization of hamster piRNAs and retrotransposons

While female hamster piRNAs have been investigated elsewhere (24), we characterized testicular piRNAs at 9 days postpartum (dpp) when spermatogonia form, at 13 dpp when testes contain spermatogonia, and at 21 dpp when spermatogenesis reaches the pachytene stage of meiosis (25). Hamster testicular piRNAs shared many features with those of other mammals. Unique and repetitive 27-29 nt “pre-pachytene” piRNAs and highly abundant non-repetitive 29-30 nt “pachytene” piRNAs had typical piRNA sequence features (Fig. 1B, S1 and S2). Pre-pachytene piRNAs were broadly dispersed at a relatively low density in intergenic and genic regions, while most pachytene piRNAs mapped to ~100 loci, many of which were syntenic with mouse, cow, and human testicular piRNA loci (Fig. S3, Table S1-S3).

Since piRNAs provide an adaptive defense against TEs, we characterized the nexus between hamster piRNAs and TEs. Using an improved golden hamster genome assembly (24), we determined the entire complement of hamster TEs, identified potentially active retrotransposon subfamilies, and estimated the abundances of retrotransposon-derived piRNAs. Analysis of long terminal repeat (LTR) retrotransposons revealed their distinct evolutionary paths from mice (Data S1-S3 and Supplementary text). While there was no notable expansion among autonomous elements from the ERVL family (Data S2 and Fig. S4A), we observed that ERVK family retrotransposons, exemplified by MYSERV and Intracisternal A Particle (IAP) elements, expanded during the hamster genome evolution (Data S1 and Fig. S4C). IAP retrotransposons evolved in hamsters and mice from an ancestral retrovirus (26,27). In the golden hamster genome, we identified 110 full-length intact (FLI) IAP insertions classified as IAPLTR3/4 (Data S4, Fig. S4C) while the IAPE subgroup is active in mice (28). Remarkably, IAP-derived piRNAs were the second most abundant TE-targeting piRNAs after MYSERV-derived piRNAs at 9 dpp (Fig. 1C and S4A). Consistent with that, we observed high abundances of pre-pachytene and pachytene piRNAs antisense to FLI IAP (Fig. 1D). Analysis of non-LTR LINE L1 elements, the most successful TEs invading mammalian genomes (29), revealed 110 FLI L1 elements from the Lx5/6 subfamily (Data S5), which is comparable to the 146 FLI L1s in the human genome but much smaller than the 492 in the rat genome (30) or the 2,811 in the mouse genome (30) (Fig. S4D). Analysis of piRNA sequences suggested that hamster FLI L1s are targeted mainly by pre-pachytene antisense piRNAs (Fig. 1E).

### Post-meiotic and post-zygotic sterile phenotype of female*Mov10l1* −/− hamsters

To examine the biological role of the hamster piRNA pathway, we knocked-out *Mov10l1* by deleting the exon 20 encoding the helicase domain (Fig. 1F and S5), generating an analogous deletion studied in mice (3). Western blot analysis showed lack of MOV10L1 in *Mov10l1*^−/−^ testes (Fig. 1G). Heterozygotes were fertile and segregation of genotypes did not deviate significantly from the Mendelian ratio but homozygotes of both sexes were sterile (Table 1). Male *Mov10l1*^−/−^ sterility could be expected, but female *Mov10l1*^−/−^ sterility was surprising given the normal fertility of female *Mov10l1*^−/−^ mice (3,4).

**Table 1.**
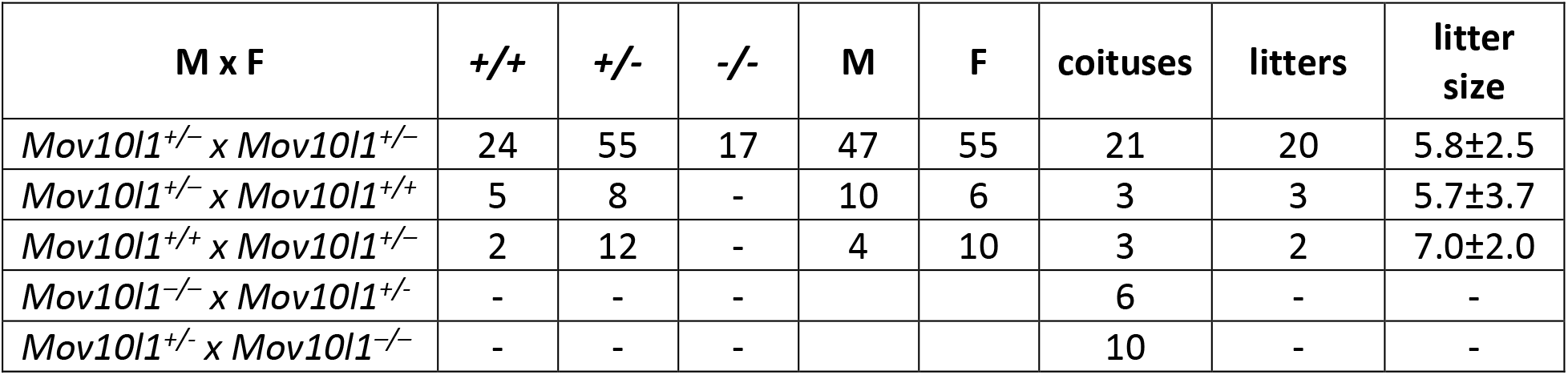
Mating performance

| **M x F** | **+/+** | **+/-** | **-/-** | **M** | **F** | **coituses** | **litters** | **litter size** |
| --- | --- | --- | --- | --- | --- | --- | --- | --- |
| *Mov10l1*<sup>+/-</sup> x *Mov10l1*<sup>+/-</sup> | 24 | 55 | 17 | 47 | 55 | 21 | 20 | 5.8±2.5 |
| *Mov10l1*<sup>+/-</sup> x *Mov10l1*<sup>+/+</sup> | 5 | 8 | - | 10 | 6 | 3 | 3 | 5.7±3.7 |
| *Mov10l1*<sup>+/+</sup> x *Mov10l1*<sup>+/-</sup> | 2 | 12 | - | 4 | 10 | 3 | 2 | 7.0±2.0 |
| *Mov10l1*<sup>-/-</sup> x *Mov10l1*<sup>+/-</sup> | - | - | - |  |  | 6 | - | - |
| *Mov10l1*<sup>+/-</sup> x *Mov10l1*<sup>-/-</sup> | - | - | - |  |  | 10 | - | - |

To understand the basis of female *Mov10l1*^−/−^ hamster infertility, we examined ovarian histology. *Mov10l1*^−/−^ ovaries appeared normal suggesting that mutant oocytes enter the first meiotic block and develop to the fully-grown germinal vesicle-intact (GV) stage in antral follicles (Fig. 2A). *Mov10l1*^−/−^ oocytes were able to mature *in vivo* to the second meiotic block at the metaphase II (MII) stage; spindle defects among *Mov10l1*^−/−^ MII eggs were not observed (Fig. 2B and S6). Ovulated MII eggs were able to be fertilized and undergo cleavage as evidenced by 1- and 2-cell zygotes obtained from mated *Mov10l1*^−/−^ females (Fig. 2C and S7). These results show *Mov10l1*^−/−^ GV oocytes retain meiotic competence but lack developmental competence. Consistent with this conclusion, female *Piwil1^−/−^* hamsters also show post-zygotic sterility and female *Piwil3^−/−^* hamsters show post-zygotically reduced fertility (31).

**Figure 2.**
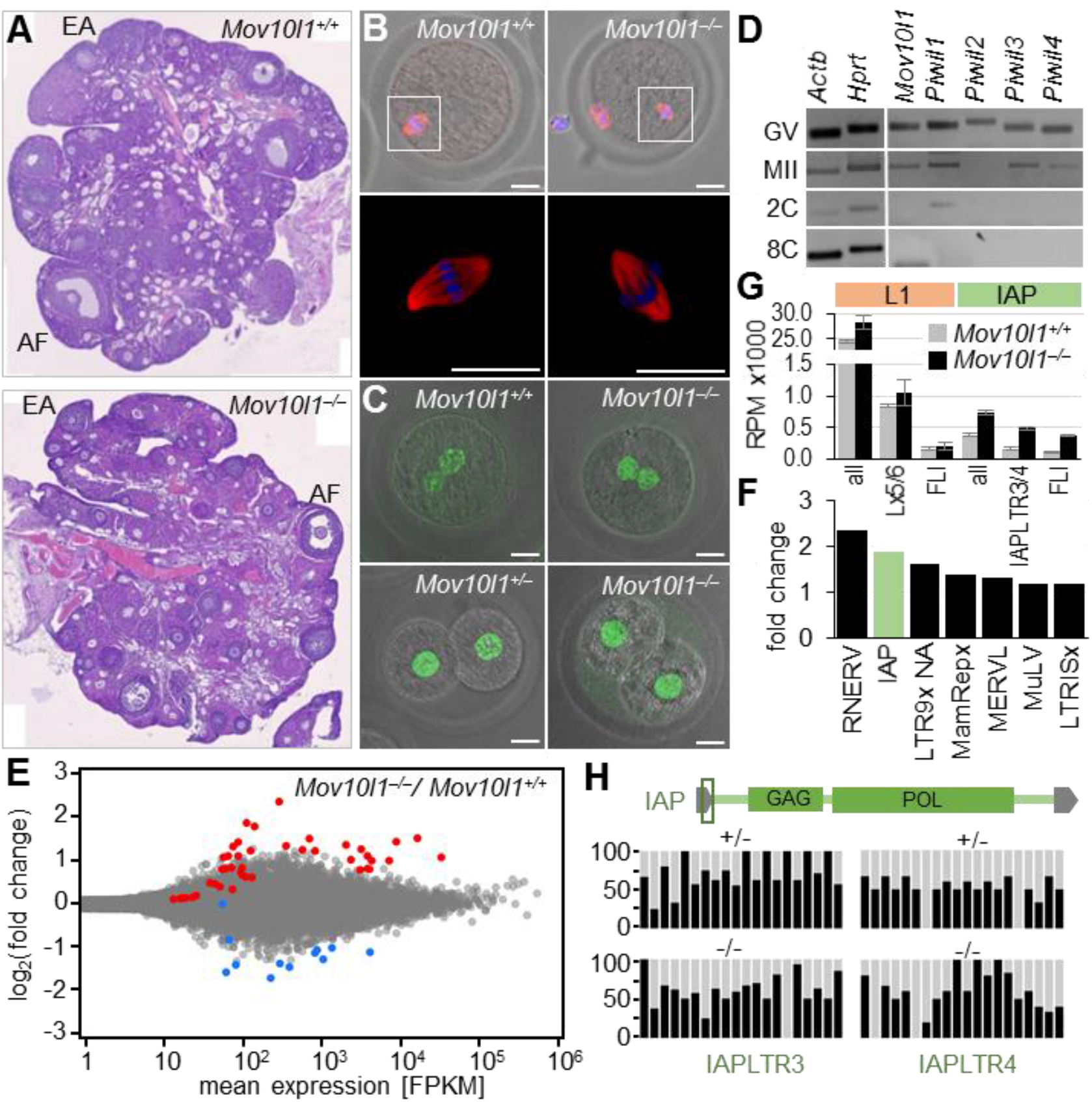
Female *Mov10l1^−/−^* phenotype. (A) Hematoxylin-eosin staining of ovarian sections. Antral follicles (AF) and early antral follicles (EA) indicate normal follicular development. (B) Mutant oocytes mature to the MII stage with a normal spindle [red, tubulin; blue, DNA (DAPI)]. Scale bar = 20 μm. (C) Mutant eggs can be fertilized and give rise to 1-cell and 2-cell zygotes. H3K9me3 staining (green) shows that zygotes from *Mov10l1^−/−^* eggs have no major heterochromatin defect. Scale bar = 20 μm. (D) RT-qPCR analysis of transcripts of piRNA pathway genes in oocytes and zygotes. (E) Transcriptome profiling of fully-grown *Mov10l1^−/−^* oocytes. An MA plot depicting differentially expressed genes in mutant oocytes relative to wild-type oocytes. Red and blue points depict protein-coding genes with significantly higher (red) or lower (blue) transcript abundance in *Mov10l1^−/−^* oocytes (DESeq2 p-value <0.01). (F) Changes in retrotransposon expression. The graph ranks the most upregulated LTR retrotransposon groups in *Mov10l1^−/−^* oocytes. (G) Changes in levels of transcript from L1 and IAP families and subfamilies. Depicted are RPMs calculated for antisense 24-31 nt RNAs mapping to L1 or IAP elements (all), active subfamilies, and only full-length intact elements (FLI). (H) DNA methylation of IAP FLI sequences. Vertical bars represent methylation (black portion) observed for the first 20 dinucleotides 5’-CpG covered by at least four sequence reads; the analyzed region corresponds to the central and 3’ part of the 5’ LTR, as indicated in the IAP scheme.

Although maternal piRNAs might contribute to the zygotic phase of the oocyte-to-embryo transition, maternal expression of *Mov10l1* and *Piwil* genes (Fig. 2D and S8) showed that the post-zygotic *Mov10l1*^−/−^ phenotype is a maternal effect. To examine this, we investigated transcriptomes of fully-grown *Mov10l1*^−/−^ GV oocytes, which is the last pre-ovulatory stage of oogenesis. We detected significantly different expression of 57 genes between *Mov10l1*^−/−^ and *Mov10l1*^+/+^ oocytes (Fig. 2E, Table S4). When considering also *Mov10l1*^+/−^ profiling, only 13 genes had significantly increased and none had decreased transcript levels in *Mov10l1*^−/−^ relative to *Mov10l1*^+/+^ or *Mov10l1*^+/−^ GV oocytes (Fig. S9). In contrast, 1612 genes showed significant changes in transcript abundance in *Piwil1*^−/−^ MII eggs (31). Apart from a piRNA-independent role of PIWIL1 (32), this difference may be because *Piwil1*^−/−^ MII eggs build up additional transcriptome changes during meiotic maturation. However, even small transcriptome changes in *Mov10l1*^−/−^ oocytes still could contribute to the observed developmental incompetence, perhaps analogously to *Ythdf2*^−/−^ oocytes, which lack developmental competence but show only minor transcriptome disturbance (33).

Developmental competence could also be affected by de-repression of TEs. RNA-seq analysis showed a limited increase in LTR retrotransposon transcripts in *Mov10l1*^−/−^ oocytes (Fig. 2F). IAP-derived sequence reads showed a ~2-fold increase for all IAP RNAs (Fig. 2F) and 3.5-fold increase for reads perfectly matching FLIs (Fig. 2G). This suggests impaired repression of FLI-IAPs in *Mov10l1*^−/−^ oocytes, without global IAP de-repression. In contrast, fertile *Mili^−/−^* mouse oocytes showed a ~7-fold increase in transcripts matching active IAP subfamilies (18). Only a ~15% increase in L1-related reads and ~25% increase in reads mapping to FLI L1 transcripts were observed in *Mov10l1*^−/−^ oocytes (Fig. 2G).

Moderate-to-low increase of retrotransposon transcripts in oocytes was accompanied by little if any global change in DNA methylation, as estimated by whole-genome bisulfite sequencing (Fig. S10). Retrotransposon families that recently expanded in the hamster genome and were associated with high amounts of piRNAs in hamster testes (Lx5, MuLV, MYSERVx, IAP, and ORR1, Fig. 1C and S4) did not show reduced DNA methylation either (Fig. S10C). Likewise, overall DNA methylation of FLI IAP elements was present, although several CpG positions in LTR may show reduced methylation frequency (Fig. 2H and S11). Because bisulfite sequencing yielded only partial coverage of the hamster genome, loss of DNA methylation at specific loci might have escaped detection, such as at loci described in *Piwil3*^−/−^ hamster oocytes (31). In addition, the observed IAP FLIs de-repression may not involve only aberrant DNA methylation as nuclear piRNA complexes can also modify histones (34). Whatever the underlying mechanisms, our observation of post-meiotic and post-zygotic sterility expand the repertoire of critical roles of the mammalian piRNA pathway in the germline cycle.

### Pre-meiotic and meiotic defects in males *Mov10l1*−/− hamsters

Adult *Mov10l1^−/−^* testes showed severe pre-meiotic and meiotic impairment of spermatogenesis. Adult *Mov10l1*^−/−^ males were sterile and had atrophic testes (Fig. 3A) as well as epididymal ducts devoid of sperm (Fig. S12A). Histology of old males (>50 weeks) revealed aspermatogenic seminiferous tubules (Fig. 3B and S12B). Approximately 3% of adult seminiferous tubules contained small clusters of cells positive for the germ cell marker DDX4 (VASA) (35) and the marker of meiotic cells Synaptonemal Complex Protein 3 (SCP3) *(36)* (Fig. 3C and S12C). These clusters were also positive for IAP GAG protein and γH2AX (Fig. 3C). De-repression of L1 retrotransposons was absent or the anti-ORF1 antibody did not cross-react with the hamster protein (Fig. S13). The clusters of rare survivors of germ cell “atresia” thus seem to develop a secondary phenotype similar to that of *Mov10l1*^−/−^ mice, in which spermatogenesis fails upon entry into meiosis (3–5).

**Figure 3.**
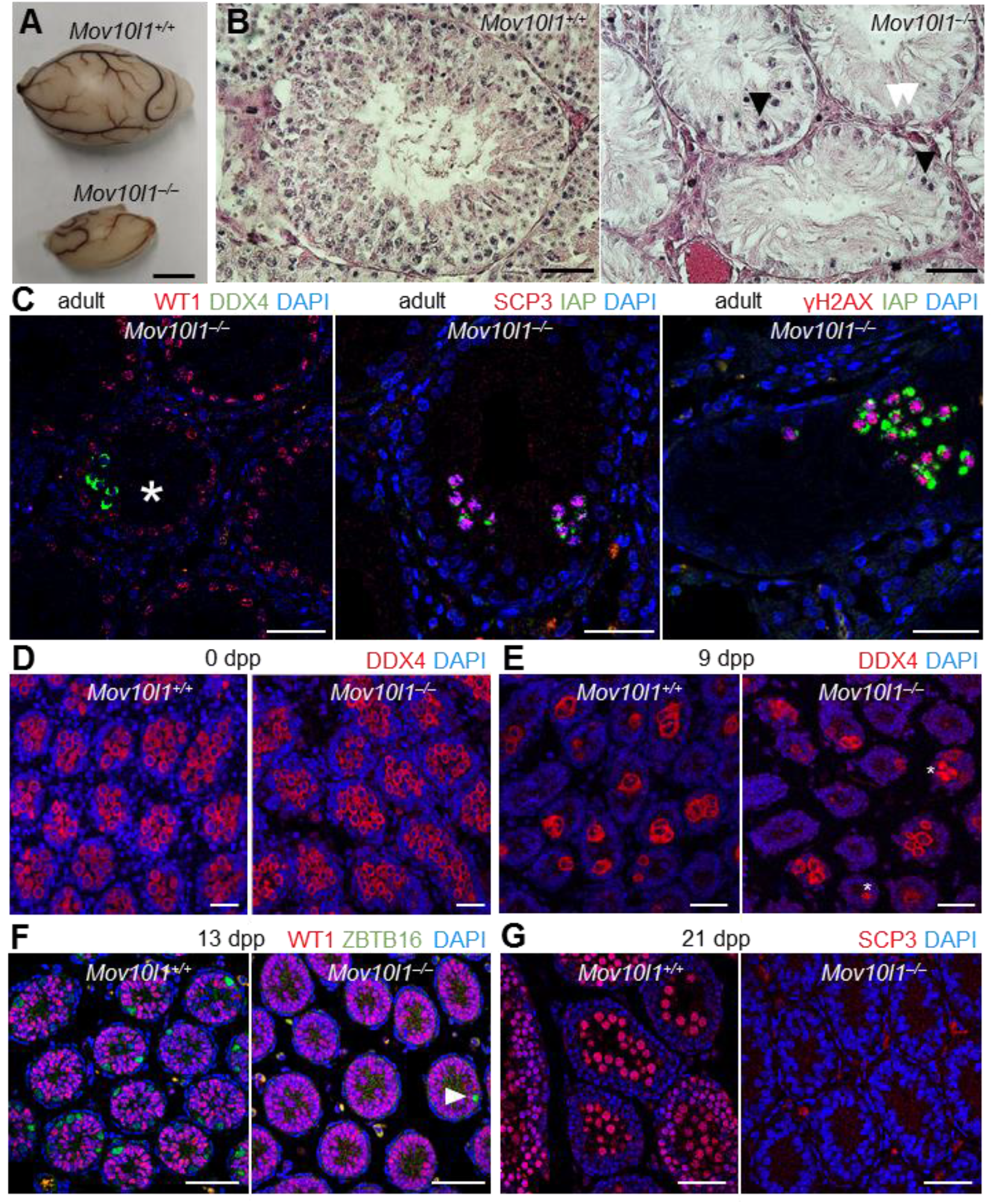
Male *Mov10l1^−/−^* phenotype. (A) *Mov10l1^−/−^* males have atrophic testes. Scale bar = 5 mm. (B) Hematoxylin-eosin staining of *Mov10l1^+/+^* and *Mov10l1^−/−^* testes. White arrowheads depict Sertoli cells, black arrowheads depict degenerated cells (see also Fig. S12). (C) Immunofluorescence staining of residual clusters of spermatogenic cells in *Mov10l1^−/−^* adult testes in order to visualize Sertoli cells (WT1) and germ cells (DDX4) (35). SCP3-positive clusters exhibit IAP expression (GAG protein, green), and DNA damage (γH2AX, red). The asterisk depicts a tubule with surviving germ cells. (D-G) Immunofluorescence staining of *Mov10l1^+/+^* and *Mov10l1^−/−^* testes at 0 dpp (D), 9 dpp (E), 13 dpp (F), and 21 dpp (G) to examine undifferentiated spermatogonia (ZBTB16) (37) and spermatocytes (SCP3) (36). DNA was stained with DAPI. Asterisks depict cells with aberrant DDX4 staining (absence of DDX4-free nucleus). The arrowhead depicts a single ZBTB16-positive spermatogonium in *Mov10l1^−/−^* testis at 13 dpp. Scale bars in panels C-G = 50 μm.

To understand the loss of germ cells in *Mov10l1*^−/−^ testes, we examined newborn (0 dpp), 9 dpp, 13 dpp, and 21 dpp animals (Fig. 3D-G). Previous work has shown newborn testes contain mitotically quiescent gonocytes, which reinitiate mitosis and are moving to the seminiferous tubule periphery by 9 dpp and give rise to spermatogonia by 13 dpp (25). In 21 dpp testes; spermatogenesis proceeds as far as the pachytene stage of meiosis I (25). We detected DDX4-positive cells in newborn and 9 dpp *Mov10l1^−/−^* testes (Fig. 3D, E). While 9 dpp *Mov10l1^−/−^* testes appeared normal, some seminiferous tubules exhibited aberrant localization of DDX4 (Fig. 3E), suggesting that the main spermatogenesis defect precedes formation of spermatogonia.

Accordingly, 13 dpp seminiferous tubules were almost devoid of cells positive for ZBTB16, the marker of undifferentiated spermatogonia (37) (Fig. 3F). At 21 dpp, we observed smaller testes, altered seminiferous tubule architecture, and complete absence of SCP3-positive meiotic cells (Fig. 3G and S14). This implied the surviving *Mov10l1*^−/−^ spermatogonia were likely compromised as they did not enter the first wave of meiosis on time. Hamster MOV10L1 is thus not necessary for embryonic development of germ cells but is required for postnatal formation of spermatogonia. The massive loss of hamster germ cells before the spermatogonia form differs from the phenotype of mice with mutations in the piRNA pathway. In *Miwi2^−/−^* mutants, spermatogonia form but are lost progressively until testes become aspermatogenic by 9 months (14,38,39).

Transcriptome profiling of 9 dpp testes identified differential expression of >900 genes (Fig. 4A and S15, Table S5). Many downregulated genes are known to be expressed in spermatogenic cells, including germline factors *Sohlh1* (40); *Ddx25* (41), *Ddx4* (35), *Dazl* (42), or many piRNA pathway components (Fig. S15B). This corroborates our hypothesis of impaired control of establishment of spermatogonia. This loss of control may have two causes. First, the loss of 26-27 nt postnatal piRNAs originating from non-repetitive sequences, including mRNA 3’ untranslated regions (8) may increase gene expression otherwise restricted by the piRNA pathway. However, we observed that only a minor fraction of upregulated gene expression appeared to be associated with higher amounts of non-repetitive piRNAs from the same loci, rather suggesting an indirect cause (Fig. S15C). Most of the observed increases in gene expression were relatively small (Fig. 4A), but expression changes may be underrepresented when genes are also expressed in other than spermatogenic cells. Notably, *Lypd6b* and *Kif5c*, two of three most upregulated genes are genomic neighbors, suggesting common regulation, which might involve *Kif5c* 3’-UTR-derived piRNAs and/or de-repression of retrotransposons in the first *Lypd6b* intron (Fig. 4B).

**Figure 4.**
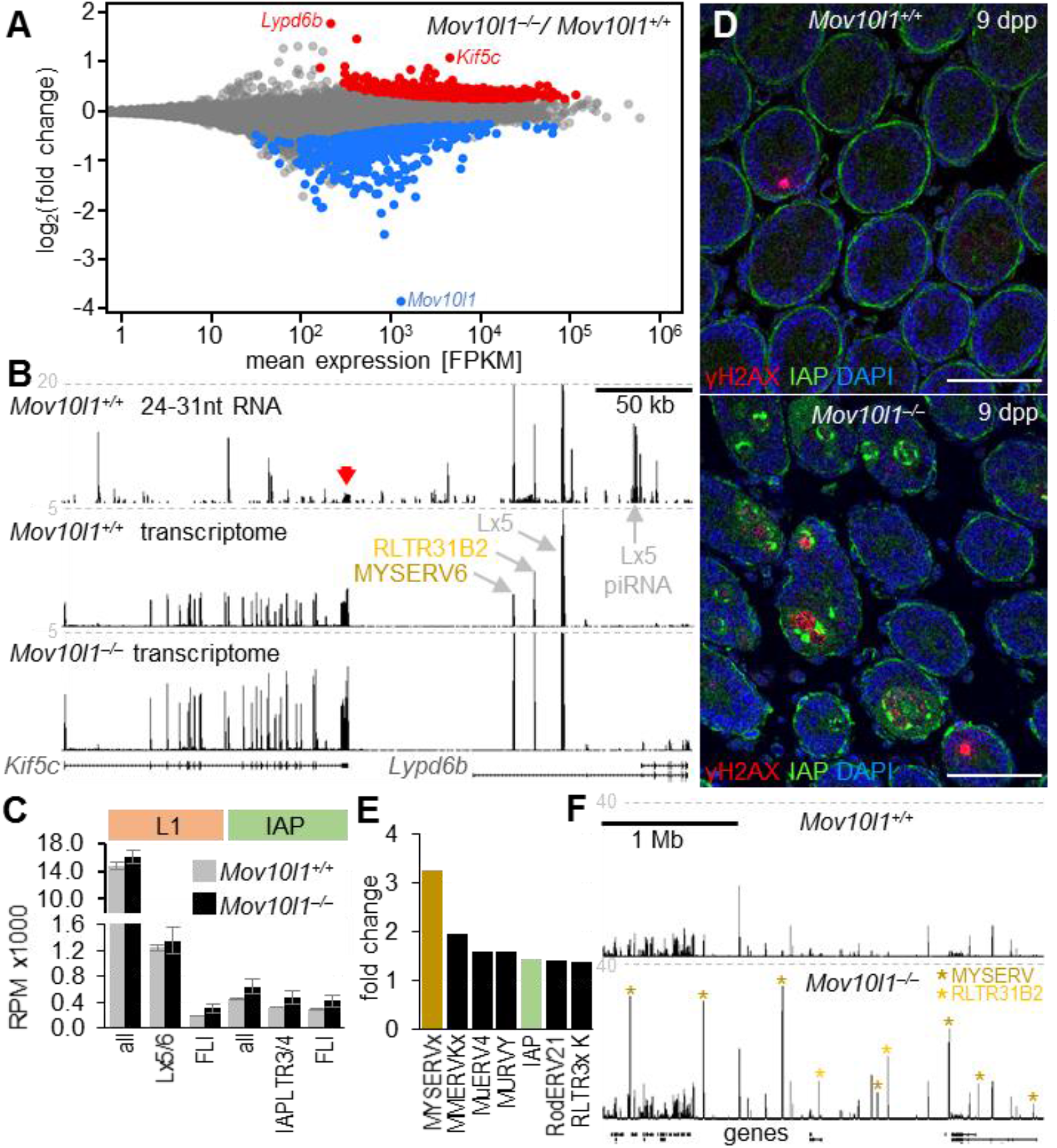
Spermatogenesis defects in *Mov10l1^−/−^* testes. (A) The MA plot depicting differentially expressed protein-coding genes: red and blue points depict genes whose transcripts were present at significantly higher (red) or lower (blue) levels in *Mov10l1^−/−^* testes (DESeq2 p-value <0.01). (B) A UCSC browser snapshot visualizing small RNAs and longer transcripts in *Mov10l1^+/+^* and *Mov10l1^−/−^* testes at the *Kif5c-Lypd6b* locus. The red arrow points to a cluster of unique piRNAs derived from the *Kif5c* 3’-UTR. Also depicted are three retrotransposon insertions in intron 1 of *Lypd6b* with increased density of mapped reads. Only perfectly mapping RNA-seq reads were used to construct the image. (C) Changes in levels of transcripts from L1 and IAP families and subfamilies. Depicted are RPMs calculated for all antisense 24-31 nt RNAs mapping to L1 or IAP elements (all), active subfamilies, and only FLI elements. (D) Immunofluorescence staining of IAP GAG (green) and γH2AX (red) in *Mov10l1^+/+^* and *Mov10l1^−/−^* testes at 9 dpp suggests IAP expression and DNA damage in germ cells inside of seminiferous tubules Scale bars = 50 μm. (E) Changes in retrotransposon expression. The graph ranks the most upregulated LTR retrotransposon groups in *Mov10l1^−/−^* testes. (F) MYSERV and related RLTR31B2 LTR-derived transcripts are upregulated in *Mov10l1^−/−^* testes at 9 dpp. A UCSC browser snapshot shows a 3-MB genomic region with upregulated retrotransposon loci (asterisks). Only perfectly mapping RNA-seq reads were used to construct the image.

In addition, de-repression of retrotransposons may also perturb gene expression at 9 dpp. Analysis of L1 and IAP FLI transcripts suggested respective approximate increases of 56% and 48% in their abundance (Fig. 4C). This mild increase contrasted with IAP and γH2AX signals in *Mov10l1*^−/−^ seminiferous tubules detected in 9 dpp but not at 0 dpp (Fig. 4D and S16). However, a similar situation concerning IAP transcript levels and immunofluorescent staining occurred when comparing mouse *G9a*^−/−^ and *Mili*^−/−^ spermatogonia (43). Notably, IAP and γH2AX patterns differed among 9 dpp seminiferous tubules. Some contained mostly IAP signal, some massive γH2AX, and some both, suggesting the onset of the phenotype occurs at 9 dpp (Fig. 4D and S16B).

Even stronger evidence that mobilization of retrotransposons may explain loss of control over spermatogenesis comes from analysis of MYSERV LTR retrotransposons. MYSERV elements give rise to highly abundant 9 dpp piRNAs (Fig. 1C and S4A). In 9 dpp *Mov10l1^−/−^* testes, MYSERV-derived reads were 3.3-fold more abundant (Fig. 4E) but many MYSERV insertion loci across the genome exhibited a ~20-fold increase (Fig. 4F). De-repression of MYSERV retrotransposons implies genome-wide failure to silence MYSERV insertion loci and may be another major, if not main, cause of the failure of germ cells to form spermatogonia. MYSERV thus seems to exemplify a hamster-specific TE threat to genome integrity, which the piRNA pathway has contained. Consequently, absence of the piRNA pathway caused a failure at a stage to which the TE had adapted its expression. Hamster-specific retrotransposon de-repression would explain why spermatogonia form in mice lacking *Mov10l1* (3–5). At the same time, the model of retrotransposon-induced failure of spermatogenesis is consistent with rapid demise of mouse spermatogonia where retrotransposon de-repression was enhanced by mutating *G9a* in addition to *Mili* (44).

Our results, complemented by those of Hasuwa et al. (31), uncover critical pre-meiotic, meiotic, and post-meiotic functions of the piRNA pathway in the golden hamster and substantially expand the known contributions of the pathway to control of the mammalian germline cycle. More importantly, substantial differences between hamster and mouse *Mov10l1^−/−^* phenotypes demonstrate the adaptive nature of the piRNA pathway, which leads to new gene regulation and flexibly protects the germline cycle against retrotransposons whenever a new threat emerges.

## MATERIALS & METHODS

### Animals

Golden (Syrian) hamsters *Mesocricetus auratus* were purchased from Japan SLC, Inc (for knock-out production and initial breeding) and from Janvier Labs, France (subsequent breeding). Animals were housed under controlled lighting conditions (daily light period 7:00 to 21:00 and 6:00 to 18:00 in Japan and Czechia, respectively) and provided with water and food *ad libitum*.

The animals used for experiments were euthanized by intraperitoneal injection of a lethal dose of Euthasol (Samohyl). All animal experiments were approved by the Animal Experimentation Committee at the RIKEN Tsukuba Institute (T2019-J004) and the Institutional Animal Use and Care Committee at the Institute of Molecular Genetics of the Czech Academy of Sciences (approvals no. 42/2016 and 70/2018) and were carried out in accordance with the law.

### Production of Mov10l1 mutants

Production of knock-out hamsters was carried out using an *in vivo* electroporation CRISPR/Cas9 system as described previously (23,45). Pairs of sgRNAs were designed to cleave *Mov10l1* genomic sequence in intron 19 (sequence of DNA targets: 5’-GGGTATCACATGACTTGGGG – 3’; 5’-GGTGTTGGGATCATAGTGGGG – 3’) and intron 20 (sequence of DNA targets: 5’-TCTCCACTCTTCCATGTGGGG-3’; 5’-TACCATTACATTTGTCAGGGG-3’) in order to delete the exon 20 (Fig. 1F).

Five animals were born of which one did not exhibit any deletion, two appeared homozygous for the deletion and were not used for breeding, and one male and one female showed modification of one allele (Fig. S5A). The male was fertile but did not transmit the allele while the female (#4) transmitted the allele into progeny (# of progeny = 10; 3 males + 4 females carried the mutant allele) when bred to a wild type animal. Subsequent breeding of heterozygotes for two generations with wild type outbred animals was performed in order to minimize possible off-targeting and inbreeding effects when heterozygotes were mated to produce homozygotes. The allele was confirmed by sequencing (Fig. S5B). RNA-seq showed loss of signal over exon 20 and strongly reduced transcript level of the mutant transcript (Fig. S5C).

For genotyping, ear biopsies were lysed in PCR friendly lysis buffer with 0.6 U/sample Proteinase K (Thermo Scientific) at 55 °C, with 900 rpm shaking until dissolved (approx. 2.5 h). Samples were heat-inactivated at 90 °C, 10 min and lysate was used for nested PCR reaction. Genotyping primers are provided in Table S6.

### Superovulation and zygote collection

Golden hamster females were induced to superovulate by intraperitoneal injection of 15 or 25 IU pregnant mare’s serum gonadotropin (PMSG, ProSpec Bio) at 10:00 h on the day of post-estrous vaginal discharge (day 1 of estrous cycle). 25 IU human chorionic gonadotropin (hCG, Sigma) was injected 76 h later (14:00 on day 4 of estrous cycle) and females were mated to fertile males at 18:00 on the same day.

Zygotes were collected from oviducts at 10:00 on the second day post-mating by flushing oviducts with M199TE medium: M199 medium (with HEPES, sodium bicarbonate and Eagl’s salts, Gibco BRL, Grand Island, NY) supplemented with 5% fetal bovine serum (Sigma) inactivated for 30 min at 56 °C, 5mM taurine, 25 μM EDTA and pre-equlibrated with 5% CO2, 5% O2 and 90% N2 at 37 °C. Zygotes were isolated in a dark room with red filters on the microscope light source.

### Oocyte collection

Fully-grown germinal vesicle-intact oocytes were collected from ovaries by puncturing antral follicles with a syringe needle in M2 medium (Sigma) containing 0.2 mM 3-isobutyl-1-methyl-xanthine (Sigma) to prevent resumption of meiosis. Ovulated unfertilized eggs arrested at methaphase II (MII) were collected from oviducts of superovulated females approximately 17h after hCG injection as described above. MII eggs were released from cumulus cells upon incubation with 0.1% bovine testes hyaluronidase (Sigma) in M199TE medium at 37°C for 1 min and washed three times in equilibrated M199TE medium kept under paraffin oil.

### Western blotting

Hamster and mouse tissues were homogenized mechanically in RIPA lysis buffer supplemented with 1x protease inhibitor cocktail set (Millipore) and loaded with SDS dye. Protein concentration was measured by Bradford assay and 60 μg of protein was used per lane. Proteins were separated on 6% polyacrylamide gel and transferred on PVDF membrane (Millipore) using semi-dry blotting. The membrane was blocked in 5% skim milk in TTBS, MOV10L1 was detected using anti-MOV10L1 primary antibody (4) (kind gift from J. Wang) diluted 1:250 and incubated overnight at 4°C. Secondary antibody HRP-anti-Rabbit (Thermo Fisher Scientific) was diluted 1:50,000 and signal was detected using Supersignal west femto substrate (Thermo Scientific). For TUBA4A detection, samples were run on 10% PAA gel and incubated with anti-Tubulin (Sigma, #T6074) mouse primary antibody diluted 1:10,000 and anti-mouse-HRP secondary antibody (Thermo Fisher Scientific) diluted 1:50,000.

### RT-PCR analyses

Oocyte and embryo expression analysis: Five to ten oocytes or embryos were collected per one sample in 3 μl of PBS and snap-frozen in liquid nitrogen, the number of oocytes/embryos were kept constant in individual sample sets. Next, the samples were lysed by mixing an equal volume of the 2x lysis buffer (46). Crude lysate was used for reverse transcription with SuperScript III (Thermo Scientific). Equal fraction of total RNA per oocyte/zygote was reverse transcribed by SuperScript III RT (Thermo Fisher Scientific) with random hexamers as recommended by the manufacturer. To avoid genomic DNA amplification, primers were designed to span multiple exons.

cDNA was amplified by ExTaqHS (TaKaRa) using the following program: 94°C 2 min; 35x 94 ℃ 10 s, 60 °C 30 s, 72 °C 30 s; Final extension 72 °C 3 min. The PCR products were resolved on 1.5% agarose gels and visualized using ethidium bromide. All PCR products were sequenced after cloning into pCR4 TOPO vector (TOPO-TA cloning kit for sequencing; Thermo Fisher Scientific). Primers are provided in Table S6.

### Histology and immunofluorescence on histological sections

Ovaries and testes were fixed in Hartman’s fixative (Sigma, #H0290) or 4% paraformaldehyde (PFA) in PBS for 1.5 h or overnight at 4°C. Tissues were dehydrated in ethanol, embedded in paraffin, sectioned to 2.5 – 6 μm thick sections and stained with hematoxylin and eosin or used for immunofluorescence staining.

For immunofluorescence staining of testes, sections were deparaffinized and then boiled for 18 min in 10 mM pH 6 sodium citrate solution for antigen retrieval. After 45 min blocking with 5% normal donkey serum and 5% bovine serum albumin (BSA) in PBS, sections were incubated for 1h at room temperature or overnight at 4 °C with primary antibody used in 1:200 dilutions: anti-LINE1 ORF1p (kindly provided by Dónal O’Carroll, University of Edinburgh), anti-SCP3 (Abcam, #ab976672), anti-ZBTB16 (Atlas antibodies, #HPA001499) and anti-ɣH2AX (Milipore, #05-636) and 1:400 dilutions: anti-DDX4 (Abcam, #ab27591 and #ab13840) and anti-WT1 (Novus Biologicals, #NB110-60011). Anti-IAP GAG antibody (kind gift of B.R. Cullen) was used in 1:500 dilution. Anti-mouse or anti-rabbit secondary antibodies conjugated with Alexa 488 or Alexa 594 (1:500; Thermo Fisher) were incubated for 1h at room temperature Nuclei were stained with 1 μg/ml DAPI for 10 min, slides were mounted in ProLong™ Diamond Antifade Mountant (Thermo Fisher) and images were acquired on DM6000 or Leica SP8 confocal microscope.

### Immunofluorescent staining of oocytes and zygotes

Oocytes and zygotes were fixed and permeabilized with 0.2% Triton X-100 in 4% paraformaldehyde for 30 min at room temperature followed by blocking in 2% BSA in PBS for 1h or kept in blocking buffer overnight. To visualize the meiotic spindle, MII eggs were stained with mouse anti-tubulin (Abcam, #ab7750) diluted 1:100 for 1h at room temperature. To visualize H3K9me3 histone modification, rabbit anti-H3K9-me3 (Upstate, #07-442) was used in 1:1000 dilution overnight at 4°C. MII eggs and zygotes were incubated with secondary antibody conjugated with Alexa 488 or Alexa 594 (Thermo Fisher) diluted at 1:500 for 1h at room temperature. DNA was stained with 1 μg/ml DAPI for 10 min. Leica SP8 confocal microscope was used for imaging.

### RNA sequencing – sequencing library preparation

For oocyte analysis, total RNA was extracted from 6 – 12 fully-grown oocytes using Arcturus Picopure RNA isolation kit (Thermo Fisher) according to the manufacturer’s protocol. RNA-seq libraries were generated using the Ovation RNA-Seq system V2 (NuGEN) followed by Ovation Ultralow Library system (DR Multiplex System, NuGEN) according to the manufacturer’s protocol. cDNA fragmentation was performed on Bioruptor sonication device (Diagenode) with 30 s ON, 30 s OFF for 18 cycles on low intensity. Libraries were amplified by 9 cycles of PCR and sequenced by 100 nt single-end reading using the Illumina NovaSeq6000 platform.

For analysis of testicular transcriptomes, total RNA was extracted from adult, 21 dpp, 13 dpp, and 9 dpp hamster testes using Sigma-Aldrich miR Premier micro RNA isolation kit according to the manufacturer’s protocol. Ribosomal RNA (rRNA) was depleted from RNA used for transcriptome analysis using Ribo-Zero rRNA Removal Kit (Human/Mouse/Rat) (Epicentre) according to the manufacturer’s protocol. rRNA depletion was confirmed by 2100 Bioanalyzer (Agilent Technologies). RNA-seq libraries were generated using NEBNext Ultra II directional RNA library Prep kit for Illumina (BioLabs, #E7765S) according to the manufacturer’s protocol. RNA-seq libraries from adult, 21 dpp and 13 dpp testes were sequenced by 150 nt pair-end reading and RNA-seq libraries from 9 dpp testes were sequenced by 75 nt single-end reading using the Illumina NextSeq500/550 platform.

For small RNA-seq analysis of testes, total RNA isolated from testes isolated as described above was used for NextFlex Small RNA-seq v3 kit (Amplicon). Libraries were prepared according to the manufacturer’s protocol with 3’ adapter ligation overnight at 16 °C, 15 cycles used for PCR amplification and NextFlex beads used for size selection. RNA-seq libraries were sequenced by 75 nt single-end reading using the Illumina NextSeq500/550 or 100 nt single-end reading using NovaSeq6000 platform.

### Bisulfite sequencing – sequencing library preparation

For bisulfite sequencing, 10 fully grown GV oocytes (an equivalent of 40 haploid genomes and 80 single-stranded DNAs after bisulfite conversion) were directly subjected to EZ DNA Methylation-Direct kit (Zymo Research) for bisulfite conversion with the following modifications: samples were digested with proteinase K at 50°C for 35 min and bisulfite conversion was performed as follows: 98°C 6 min, 64℃ 30 min, 95°C 1 min, 64°C 90 min, 95°C 1 min, 64°C 90 min. DNA libraries were prepared using EpiNext Post-Bisulfite DNA library Preparation kit according to the manufacturer’s protocol with 22 PCR cycles used for amplification. Final DNA libraries were sequenced by 250 nt pair-end sequenced on the Illumina NovaSeq6000 platform.

### Bioinformatic analyses

The code used for sequencing data analysis is available through GitHub: https://github.com/fhorvat/bioinfo_repo/tree/master/papers/piRNA_2021.

The list of analyzed sequencing libraries is provided in Table S7, Raw data were deposited in the GEO database as GSE164658.

#### RNA-seq and differential gene expression analysis

Raw RNA-seq reads were mapped to mouse (mm10), human (hg38), cow (bosTau9), rat (rn6), golden hamster (mesAur1) and the newest golden hamster (PRJDB10770 (24)) genomes using STAR 2.7.3a (47) with following parameters:

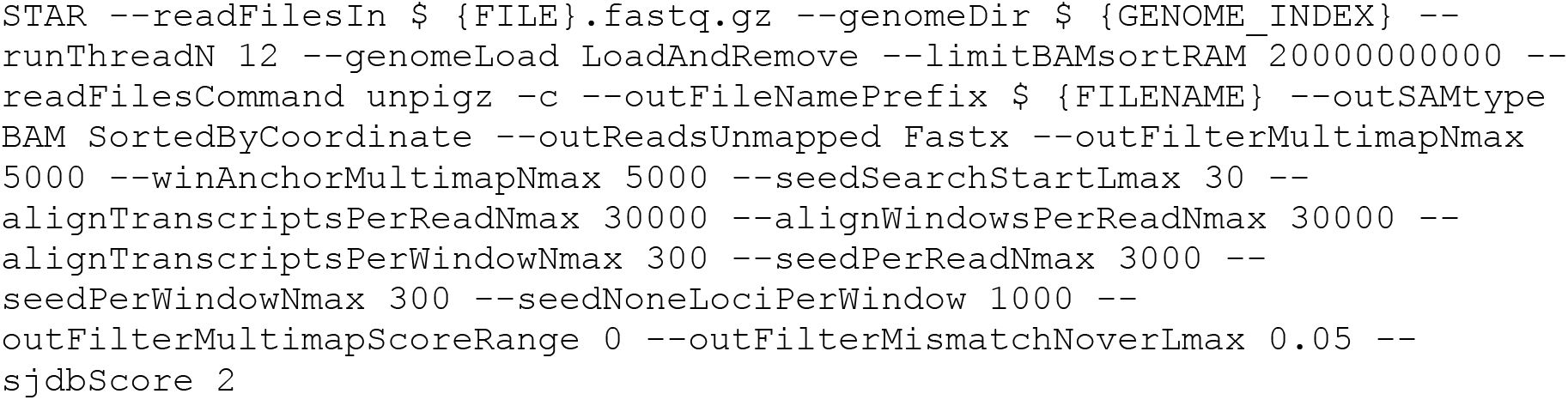

Those parameters were chosen to optimize mapping for quantification of transposable elements (48).

For analysis of expression of protein coding genes, reads were mapped with maximum of 20 multimapping alignments allowed. Reads mapped to mesAur1 were counted over exon features annotated by Ensembl (release 99) using featureCounts v2.0.0 (49):

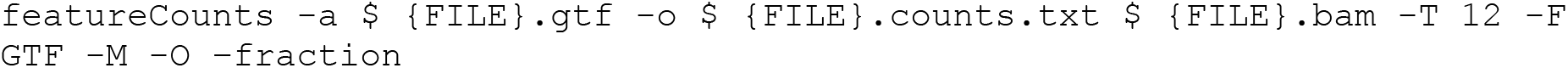

For the paired-end libraries -p flag was added. For the stranded libraries, -s 2 flag was added. Statistical significance and fold changes in gene expression were computed in R using DESeq2 package (50). Genes were considered to be significantly up-or down-regulated if their corresponding p-adjusted values were smaller than 0.01. Principal component analysis was computed on count data transformed using regularized logarithm (rlog) function.

For the heatmap showing expression of piRNA factors in testes and oocytes of four mammalian species (Fig. 1A), following publicly available datasets were used: bovine oocyte GSE52415 (51), bovine testis PRJNA471564 (52), human oocyte GSE72379 (53), human testis GSE74896 (54), mouse oocyte GSE116771 (55), mouse testis GSE49417 (56), rat oocyte GSE137563 (57), rat testis GSE53960 (58). Read mapping coverage was visualized in the UCSC Genome Browser by constructing bigWig tracks using the UCSC tools (59)

#### Small RNA-seq analysis

Small RNA-seq reads were trimmed in two rounds using bbduk.sh version 38.87 (https://jgi.doe.gov/data-and-tools/bbtools/). First, NEXTflex adapter were trimmed from right end:

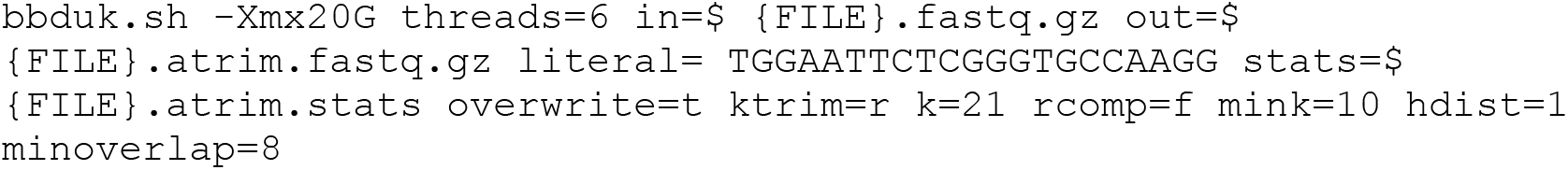

Next, 4 random bases from both sides of reads were trimmed:

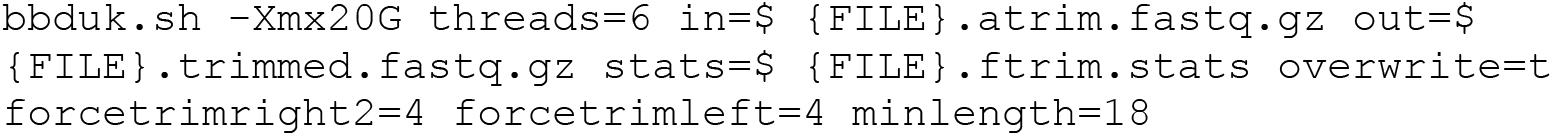

Trimmed reads were mapped to the genomes with following parameters:

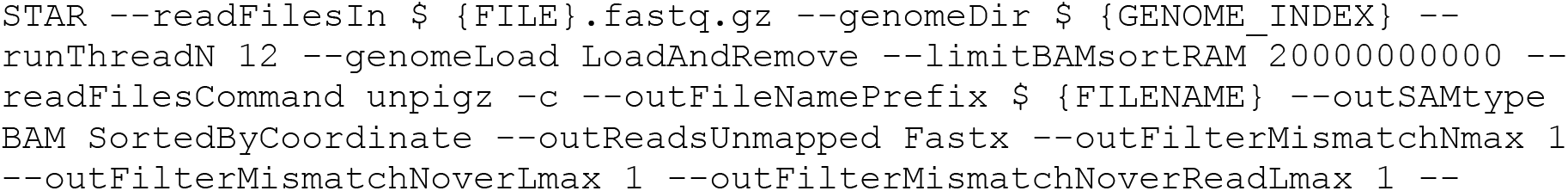
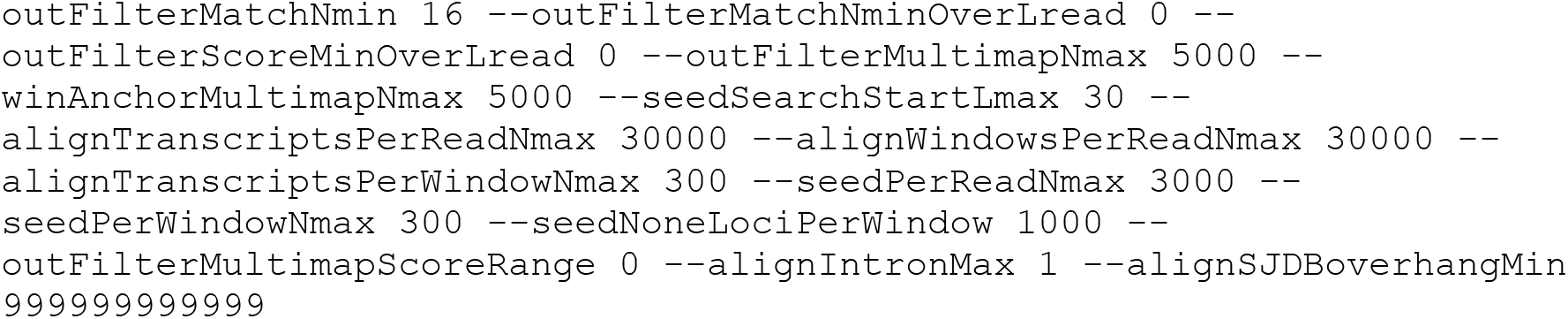

#### Definition of hamster piRNA clusters and their analysis

Small RNA sequencing (RNA-seq) analysis of whole testes was used to distinguish pre-pachytene pre-meiotic piRNAs and meiotic pachytene piRNAs as 9 dpp and 13 dpp testes contain only pre-meiotic spermatogonia while spermatogenesis in pubertal hamsters reaches the pachytene stage at 21 dpp (25). piRNA clusters in testes were defined using custom R scripts as follows:

1. Genome was divided into 1 kb windows and for each window alignments of 24-31nt reads were counted with fractional counts. Counts were normalized with total number of 19-32 nt reads in millions into RPM. RPMs were then normalized for length of windows without counting gaps in assembly (N nucleotides) into RPKM values. Windows with RPKM < 1 were removed.
2. For pre-pachytene clusters, neighboring tiles were merged into clusters if their log2 fold changes of KO/WT RPKMs in 9 dpp and 13 dpp were lower than −2. For pachytene clusters, neighboring tiles were merged into clusters if their log2 fold changes of KO/WT RPKMs in 21 dpp were lower than −2 and log2 fold changes of KO/WT RPKMs in 13 dpp were higher than −2.
3. Next, clusters were merged into super-clusters if they were at most 2kb apart.
4. Clusters were manually curated and final RPKM values were recalculated.

For further analysis, we selected loci with piRNA density above 10 reads per million (RPM) per kilobase for 9 dpp 13 dpp pre-pachytene piRNAs and 100 rpm for 21 dpp pachytene piRNAs (Fig. S3A).

Pre-pachytene piRNAs in 9 dpp and 13 dpp testes were much less abundant, relative to both, miRNAs and 21 dpp pachytene piRNAs (Fig. S1). They were also dispersed across the genome in many more intergenic and genic regions but at much lower density than observed for pachytene piRNAs (Fig. S3, Table S1 and S2). Majority of pre-pachytene piRNAs started with U. A at the position 10 (10A) was observed among longer piRNAs (29 nt) suggesting that a fraction of piRNAs originates through ping-pong mechanism (Fig. S2).

Consistent with mouse data (60), pachytene piRNAs dominated the small RNA population at 21 dpp, their length peaked at 30 nt (Fig. S1), most of them originated from a limited (~100) number of typically intergenic loci, and were absent at 9 dpp and 13 dpp. For further analysis we used 116 clusters with >100RPM (Table S3). When ranked by the contribution to the piRNA pool at 21 dpp, the first 20 clusters contributed 53% of 24-31 nt small RNAs in 21 dpp testes and were syntenic with piRNA-producing loci in mouse, bovine and human testes (Table S3). The first 50 clusters produced majority (>90%) of 24-31 nt small RNAs in 21 dpp testes and >90% were syntenic piRNA clusters in the mouse genome (Table S3). Most of pachytene piRNAs started with U, a small fraction of pachytene piRNAs carried non-templated U additions at the 3’ end (Fig. S2). 10A, the signature of the “ping-pong mechanism”, was weak if any in pachytene piRNAs (Fig. S2).

#### piRNA sequence logos

The sequence logos (Fig. S2) were calculated from the primary alignments only. First, only the reads with aligned first nucleotide were selected (all reads with clipped 5’-end were removed). Then, reads mapped within the pi-RNA clusters were selected. The 25-31 nt long reads were used for drawing the sequence logo (61).

#### Annotation of transposable elements

Transposable elements in new golden hamster assembly were annotated with RepeatMasker 4.0.9 (62) using mouse repeats database as closest available annotated organism.

In summarizing expression of transposable element groups, each read was counted only once using a custom R script.

#### Bisulfite sequencing

Raw bisulfite sequencing reads were trimmed with following parameters:

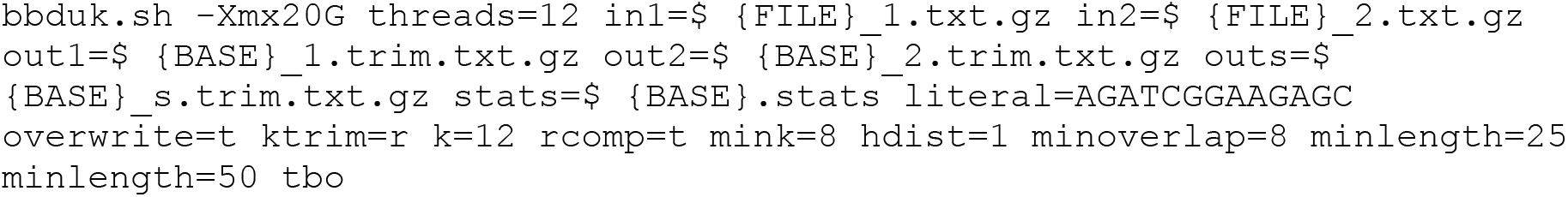

Trimmed reads were mapped to the genome using Bismark (63):

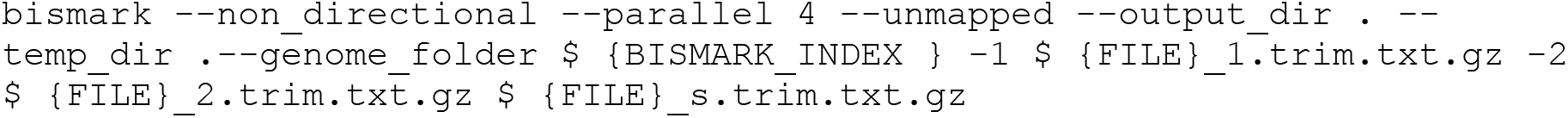

Next, in order to remove alignments arising from excessive PCR amplification, alignments were deduplicated using deduplicate_bismark script. Methylation information for individual cytosines were then extracted with bismark_methylation_extractor. For the expression of FLI retrotransposon consensus, reads were first mapped to the genome. Next, reads mapping to individual FLI insertions were extracted and mapped again to FLI element consensus sequences.

## Supporting information

Table S19 dpp pre-pachytene piRNA clusters

Table S213 dpp pre-pachytene piRNA clusters

Table S321 dpp pachytene piRNA clusters

Table S4DEG genes in Mov10l1-/- GV oocytes

Table S5DEG genes in Mov10l1-/- 9 dpp testes

Data S1ERVK LTR retrotransposon nucleotide exchange rates

Data S2Other LTR retrotransposon nucleotide exchange rates

Data S3MaLR LTR retrotransposon nucleotide exchange rates

Data S4Golden hamster IAP FLI sequences

Data S5Golden hamster LINE L1 FLI sequences

## ACKNOWLEDGMENTS

We thank Haruhiko Siomi for access to the refined golden hamster genome assembly, Dónal O’Carroll and Chapin Rodriquez for critical comments on the manuscript, Romana Šustrová for taking care of hamsters, and Šárka Kocourková (Laboratory of Genomics and Bioinformatics, IMG) for assisting with RNA-seq.

## AUTHOR CONTRIBUTIONS

ZL, HF, AO, and PS designed the experiments. HF, MH, AO generated mutant hamsters. ZL and HF analyzed the phenotype. ZL conducted RNA-seq and bisulfite-seq experiments. FH and JP performed bioinformatics analyses. ZL, HF, FH, JP, RM, and PS analyzed phenotype data. PS lead the project and drafted the first version of the manuscript, which all authors revised.

## SUPPLEMENTARY MATERIAL

## SUPPLEMENTARY TEXT

### Hamster retrotransposon analysis

As one of the key roles of piRNAs is repression of retrotransposons, we determined the hamster retrotransposon complement in order to reveal potentially active retrotransposons, their expression, and abundance of sense and antisense piRNAs carrying their sequences. Although some information could be extracted from the published *Mesocricetus auratus* Mesaur1.0 and *Cricetulus griseus* (Chinese hamster) criGriChoV2 genomes, this analysis was fragmentary owing to the incompleteness of the golden hamster genome and sequence divergence of the Chinese hamster genome. For analysis of solo long-terminal repeats (LTRs) and other short retrotransposon insertions, Mesaur 1.0 could yield acceptable results (22) but for long autonomous retrotransposon analysis, it was inadequate. For example, Mesaur1.0 contains several larger fragments of IAP retrotransposon covering most of its internal sequence but not a single full-length element matching the published full-length IAP sequence (64). L1 retrotransposon insert assemblies are even worse in Mesaur1.0. Although this could be partially remedied by using Chinese hamster genome data and raw sequencing data from Mesaur 1.0, the situation was problematic for rigorous analysis of golden hamster retrotransposons.

The issue was solved by authors of the accompanying article (31) who re-sequenced and re-assembled the golden hamster genome to a quality, which enabled a rigorous analysis (24). Subsequently, we annotated retrotransposons using the Repeatmasker and de-novo RepeatModeler. Good concordance between the two annotations suggested that no element group was missed by Repeatmasker. We subsequently followed RepeatMasker annotation to maintain consistence with annotated murine TE groups.

### LTR elements in the hamster genome

Analysis of full-length intact LTR retrotransposons included inspection of nucleotide substitution rates of over hundreds of different repeats from ERV1, ERVK, ERVL, Gypsy families (Data S1-S3), from which we selected for further analysis a representative set of LTR retrotransposons of various age and type (Fig. S4). Among LTR retrotransposons, the ERVK class appeared as the most recently expanded retrotransposon class when considering nucleotide substitution rate in retrotransposons insertions (Data S1 and S4A). Within the IAP retrotransposon family, which exhibited the lowest substitution rate, IAPLTR3 and IAPLTR4 subfamilies of IAP sequences exhibited minimal nucleotide exchange rates (Fig. S4C) suggesting very recent or ongoing retrotransposition. Indeed, we identified 110 IAP insertions with largest intact ORFs (Data S4). These IAPs are related to mouse IAPs through a common ancestral retrovirus (26), their expansion in the hamster genome is independent from mice (64). The mouse genome had a recent IAP expansion (27,65) and IAP is currently the only active autonomous LTR element in the mouse genome (28). Regarding other LTR classes, two ERV1 family element types (LTRIS and MuLV) showed low substitution rates suggesting recent or potential ongoing expansion (Data S2). MuLV has also been reported to expand in the mouse lineage relatively recently (66). Notably, there was no support for ongoing or recent expansion of autonomous ERVL elements in the hamster genome (Data S2 and Fig. 4A). This was consistent with higher substitution rates of MaLR family of non-autonomous LTR elements, whose mobility requires functional autonomous ERVL elements ((Data S3 and Fig. S4B). This contrasts with evolution of the mouse genome, where two distinct amplification bursts of MuERV-L retrotransposon occurred around 10 and 2 MYA (67). MuERV-L is transcriptionally activated during the mouse zygotic genome activation (ZGA) and its LTRs serve as a ZGA enhancer and/or alternative promoter (22,68,69). MuERV-L expansion in the mouse genome presumably mobilized MaLR and gave rise to large MTA and ORR1A subfamilies, which have low substitution rates in the mouse genome (22) but are absent in the golden hamster genome. Consequently, the golden hamster genome and its genes did not undergo such a massive rewiring and remodeling by long terminal repeats (LTRs), as the mouse germline (22,70).

### L1 elements in the hamster genome

Among autonomous mobile elements invading mammalian genomes is the most successful L1 element (reviewed in (29)), which represents non-LTR Long Interspersed Element (LINE) class. We identified L1ME3C, Lx2, Lx4B, and Lx5 as the most recently expanded L1 groups in the golden hamster genome (Fig. S4D). Among these, we discovered 110 full-length intact elements that belonged to the Lx5 L1 subfamily (Data S5). The total number of intact L1s in the golden hamster genome is very similar to the number of L1s in the human genome (71) and much smaller than the number of putatively active full-length intact L1 insertions in the rat (492) and mouse (2811) genomes (30), which represent different recently amplified subfamilies according to low nucleotide substitution rates found in a random set of insertion sequences (Fig S4D). Analysis of criGriChoV2 revealed 103 of full-length intact L1 insertions but most of them were annotated L1_1 and only 11 as Lx6 suggesting that a different closely-related L1 subfamilies expanded in both hamster species.

Taken together, *M. auratus* and *M. musculus* genomes carry full-length intact copies of L1 and IAP retrotransposons but these retrotransposon populations represent independent retrotransposon expansion events. Full-length intact L1 load in the hamster genome is reminiscent of that in the human genome and ten times lower than that in the mouse genome. The hamster genome was not subjected to a recent ERVL class expansion, which is observed in the mouse genome but exhibits some recent mobilization of the ERV1 class elements MuLV and LTRIS.

## SUPPLEMENTARY FIGURES

**Figure S1.**
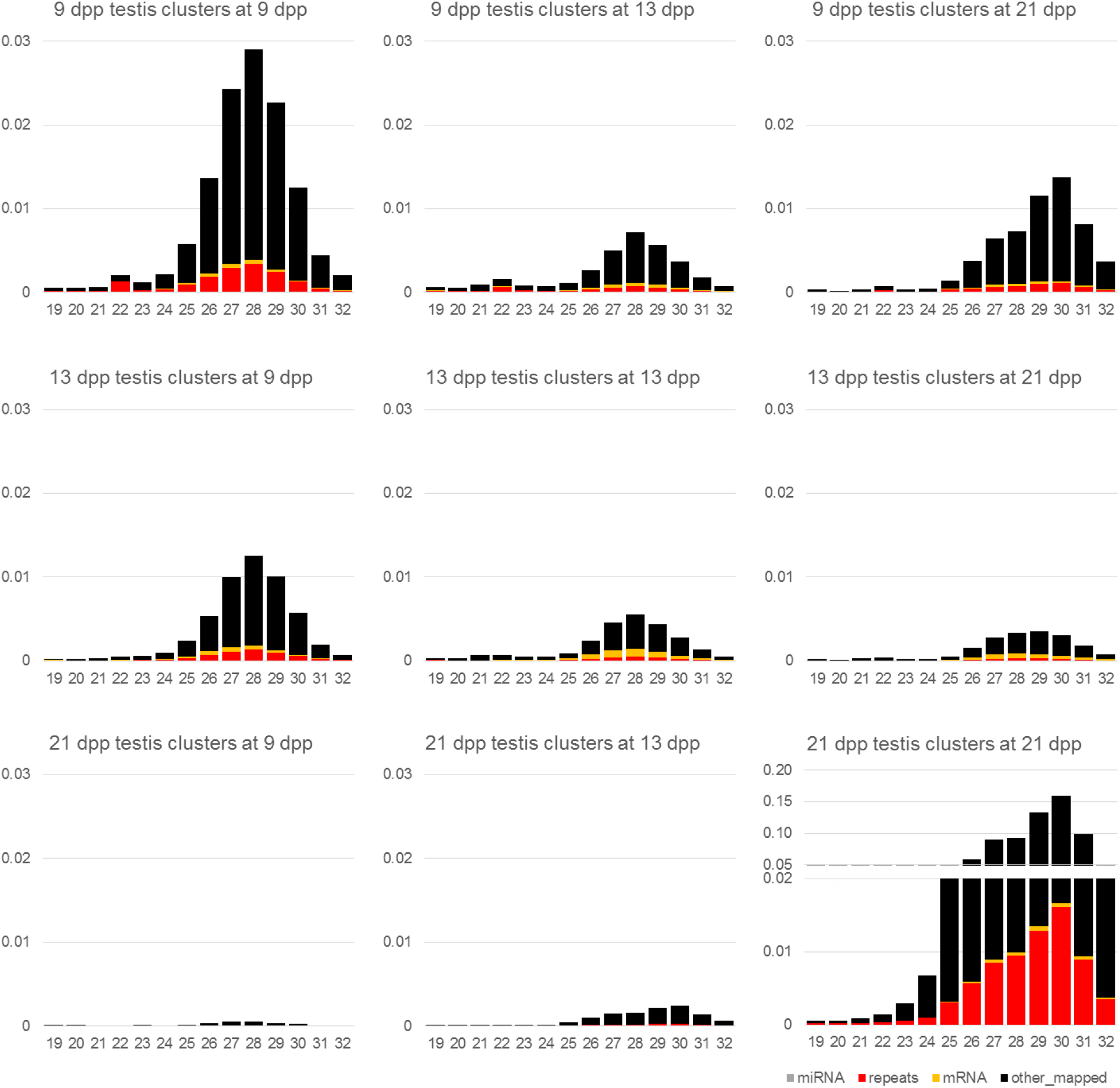
Analysis of golden hamster’s piRNA clusters I. Graphs depict frequencies of 19-32 nucleotides-long RNAs mapping to annotated pre-pachytene (9 dpp, 13 dpp) and pachytene (21 dpp) clusters (Tables S1-S3) at the three studied time points. Coloring of bars indicates proportions of miRNAs and small RNAs derived from repetitive sequences, mRNA, and other sequences.

**Figure S2.**
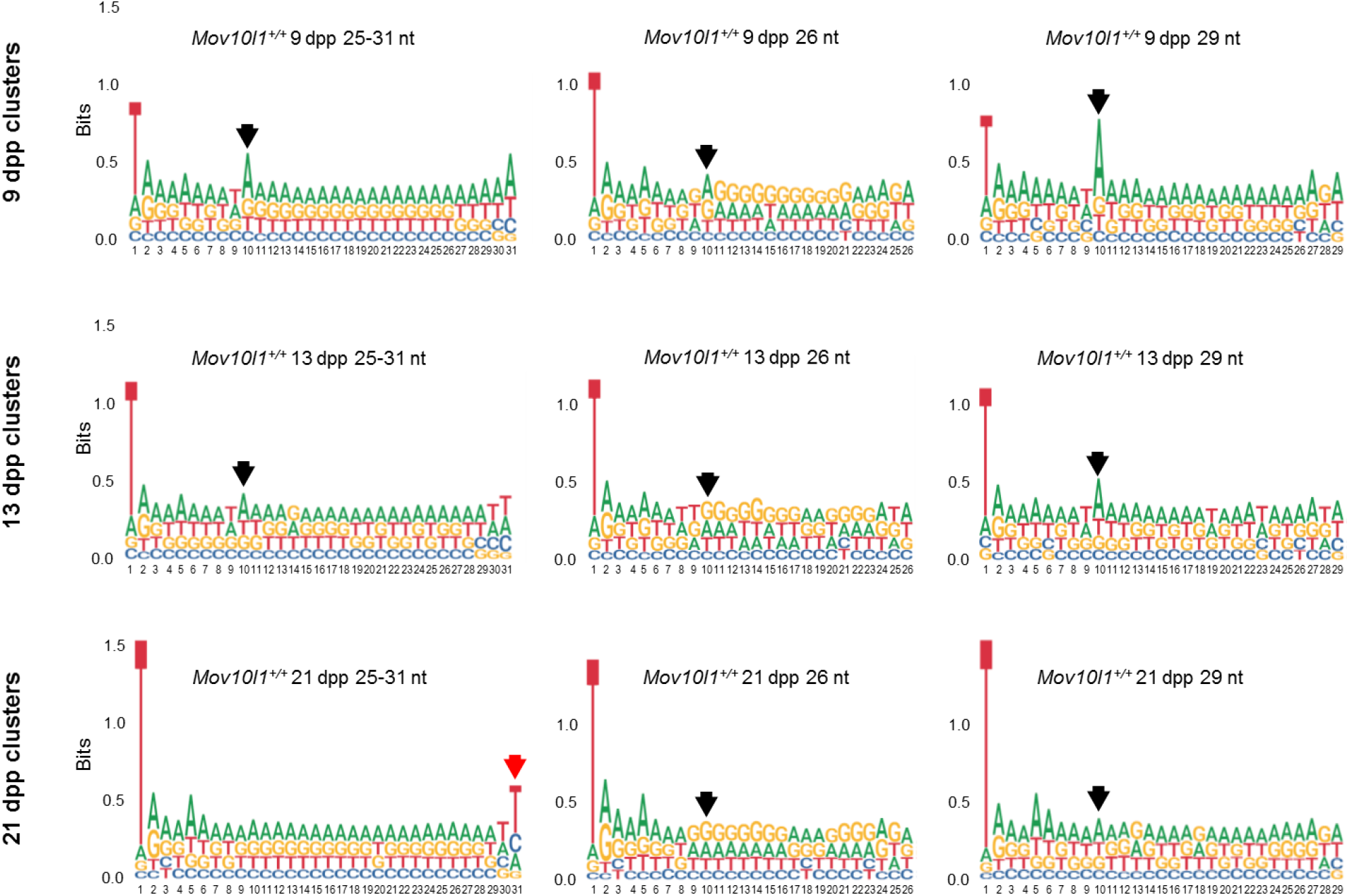
Analysis of golden hamster’s piRNA clusters II. Sequence logos for piRNA clusters from 9, 13, and 21 dpp testes (Tables S1-S3). U at position 1 is a common piRNA feature (2), preference of A at the position 10 in the sequence logos of 29 nt long piRNA clusters from 9 and 13 dpp testes is a “ping-pong signature” of secondary piRNAs (2), depicted by a black arrow. 29 nt long piRNAs from 21 dpp and shorter piRNA clusters from all three time points show little if any signs of the ping-pong signature. Red arrow points to apparent U-tailing of pachytene piRNAs. Cluster classification is the same as in Fig S1 and S3.

**Figure S3.**
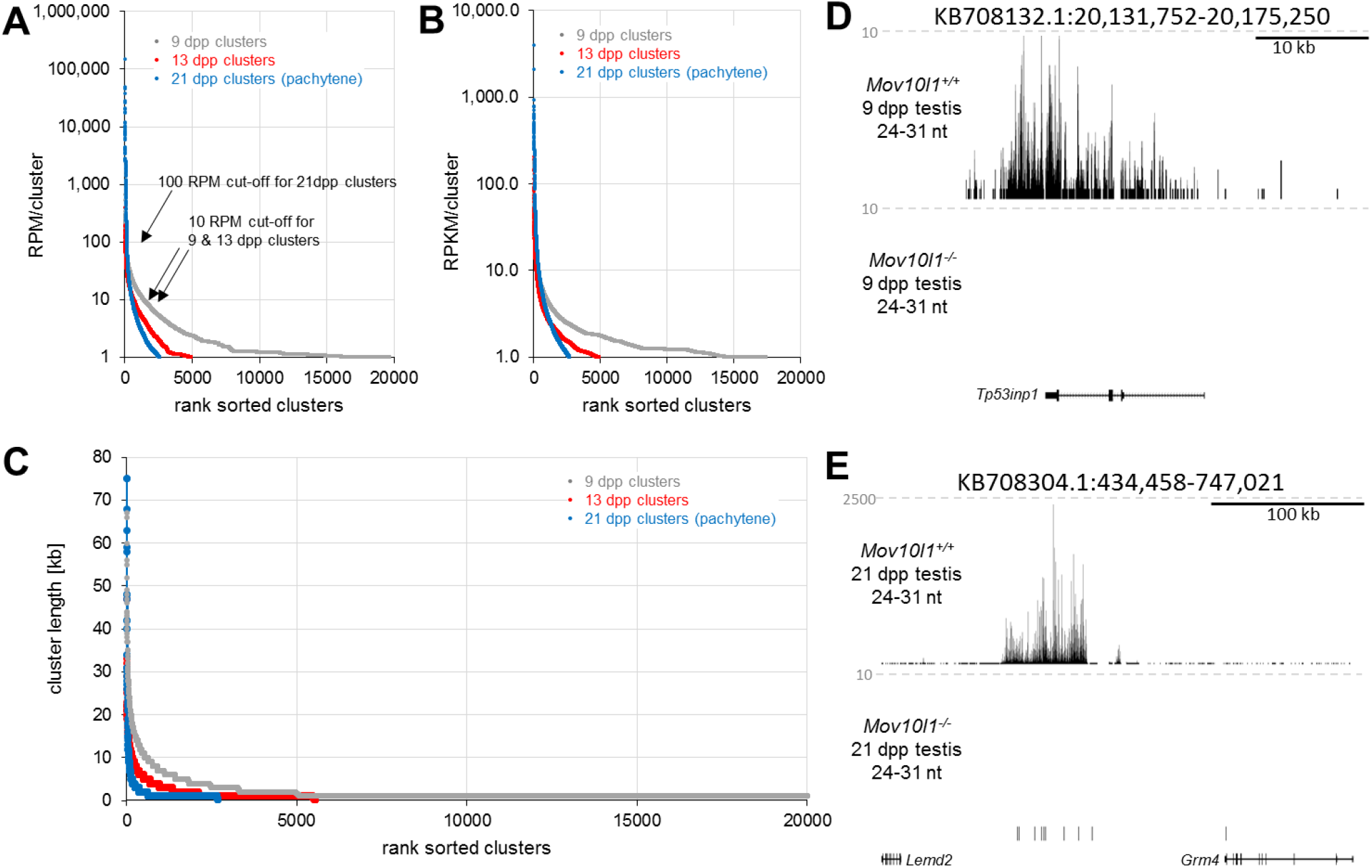
Analysis of golden hamster’s piRNA clusters III. (A) Rank-sorted annotated pre-pachytene and pachytene clusters according to the amount of small RNAs per cluster (RPM). (B) Distribution of piRNA clusters according to average piRNA density (cluster per RPKM). (C) Distribution of piRNA cluster sizes. (D) An example of a “pre- pachytene”cluster. This UCSC snapshot of 24-31 nt small RNAs mapped to an annotated 9 dpp piRNA cluster. Grey numbers indicate the scale in counts per million (CPM) 19-32 nt reads. Small RNAs from the region are absent in 9 dpp *Mov10l1*^−/−^ testes despite DDX4-positive cells are still present at this stage. (E) An example of a “pachytene” cluster. The UCSC snapshot shows extreme abundance of 24-31 nt RNAs mapping to pachytene clusters in normal testes and restriction of their presence to 21 dpp *Mov10l1*^+/+^ germ cells (*Mov10l1*^−/−^ testes at 21 dpp contain mostly somatic cells but minimum spermatogonia and no SCP3 positive cells as shown in Fig. S14).

**Figure S4.**
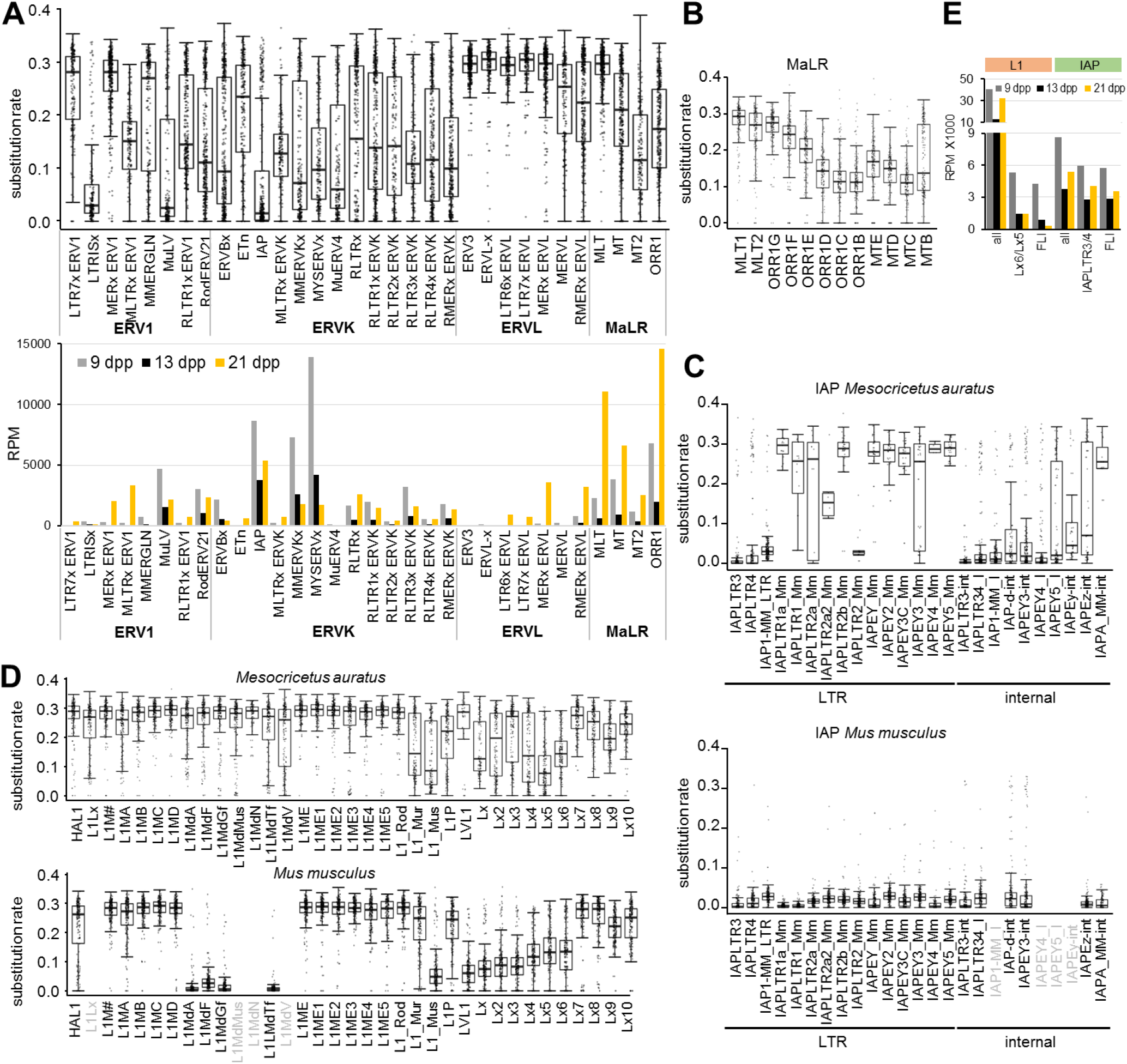
Nucleotide substitution rates of retrotransposons sequences. (A) Nucleotide substitution rates in selected LTR subgroups and abundance of 24-31 nt RNAs carrying depicted retrotransposon sequences. The lower graph depicts RPMs for 24-31 nt small RNAs per million 19-32 nt reads from wild type 9 dpp, 13 dpp and 21 dpp whole testes small RNA sequencing (B) Nucleotide substitution rates in MaLR elements. (C) Nucleotide substitution rates in IAP elements in golden hamster and mouse. (D) Nucleotide substitution rates in L1 subgroups in golden hamster and mouse. (E) Abundance of putative piRNAs from L1 and IAP sequences. All, reads from all annotated L1 or IAP sequences; Lx6/Lx5 matching subfamilies Lx5 and Lx6; IAPLTR3/4, matching IAP internal sequences of IAPLTR3 or IAPLTR4; FLI, matching full-length intact elements. y-axis represents RPMs for 24-31 nt small RNAs per million 19-32 nt reads from wild type 9 dpp, 13 dpp and 21 dpp whole testes small RNA sequencing.

**Figure S5.**
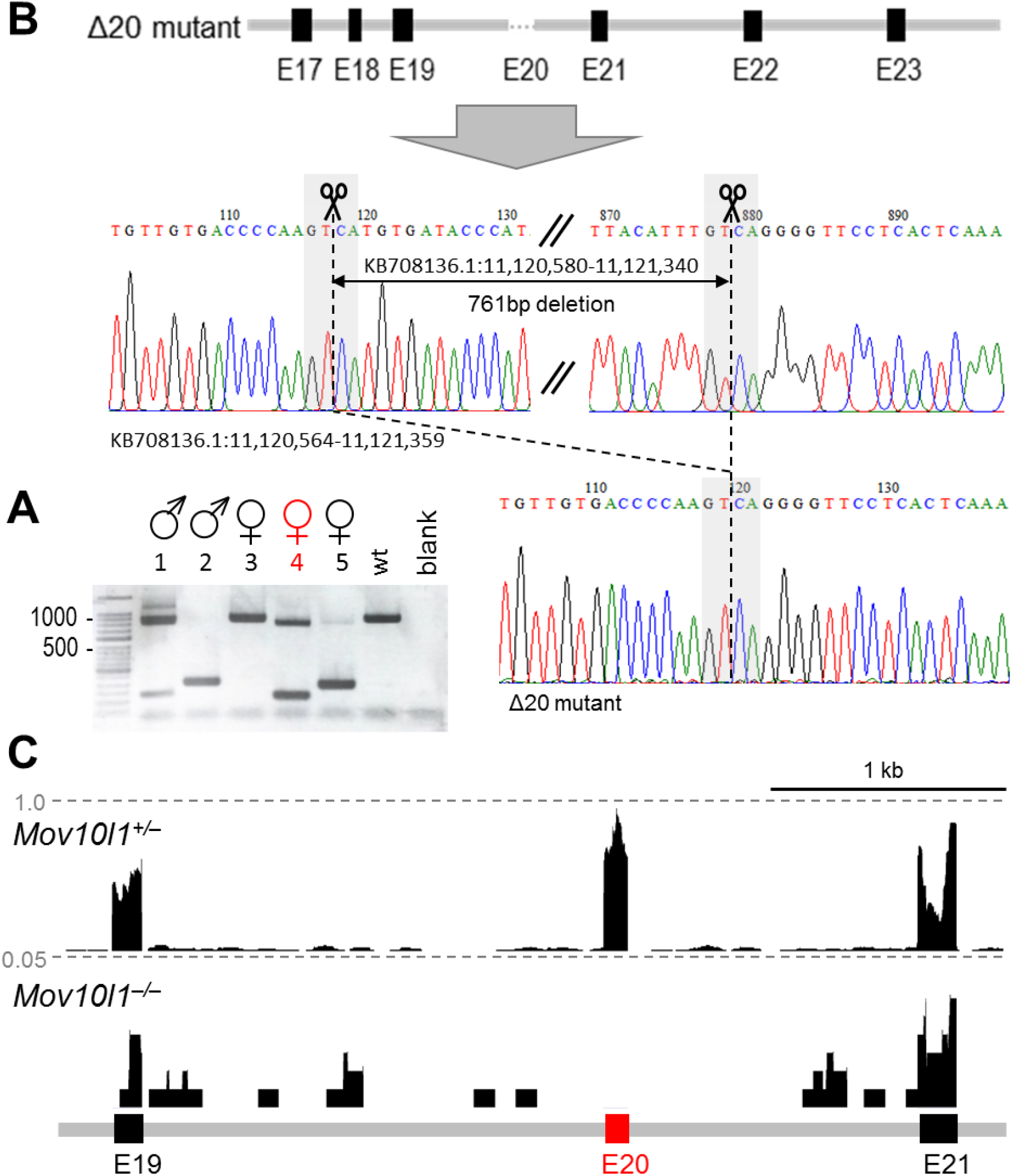
*Mov10l1^−/−^* mouse model production. (A) PCR genotyping of five F0 animals. The upper band >1 kb corresponds to the wild-type allele. 4/5 animals carried at least one mutated allele. Female #4 was heterozygous and transmitted the mutated allele into F1. (B) Schematic position of CRISPR-Cas9 cleavage sites and validation of the deletion induced in female #4 by Sanger sequencing. (C) UCSC browser snapshot of RNA-sequencing data (9 dpp testes) showing absence of *Mov10l1* sequences mapping to the removed exon 20. Dashed lines depict counts per million (CPM) of normalized (per library size) expression data.

**Figure S6.**
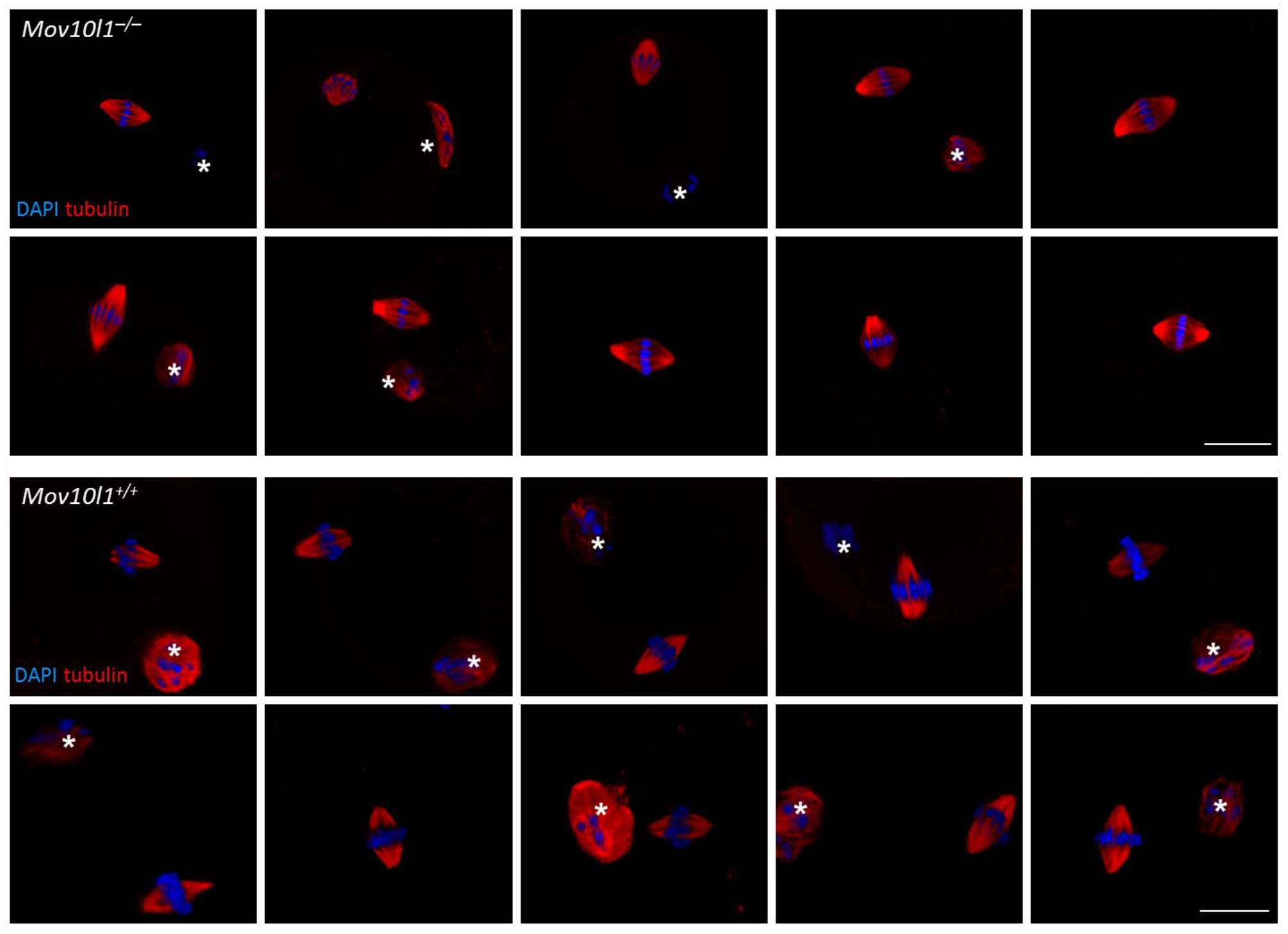
MII spindle analysis of *Mov10l1^+/+^* and *Mov10l1^−/−^* MII eggs suggests normal meiotic maturation of *Mov10l1^−/−^* oocytes. Asterisks indicate polar bodies. Scale bar = 10 μm.

**Figure S7.**
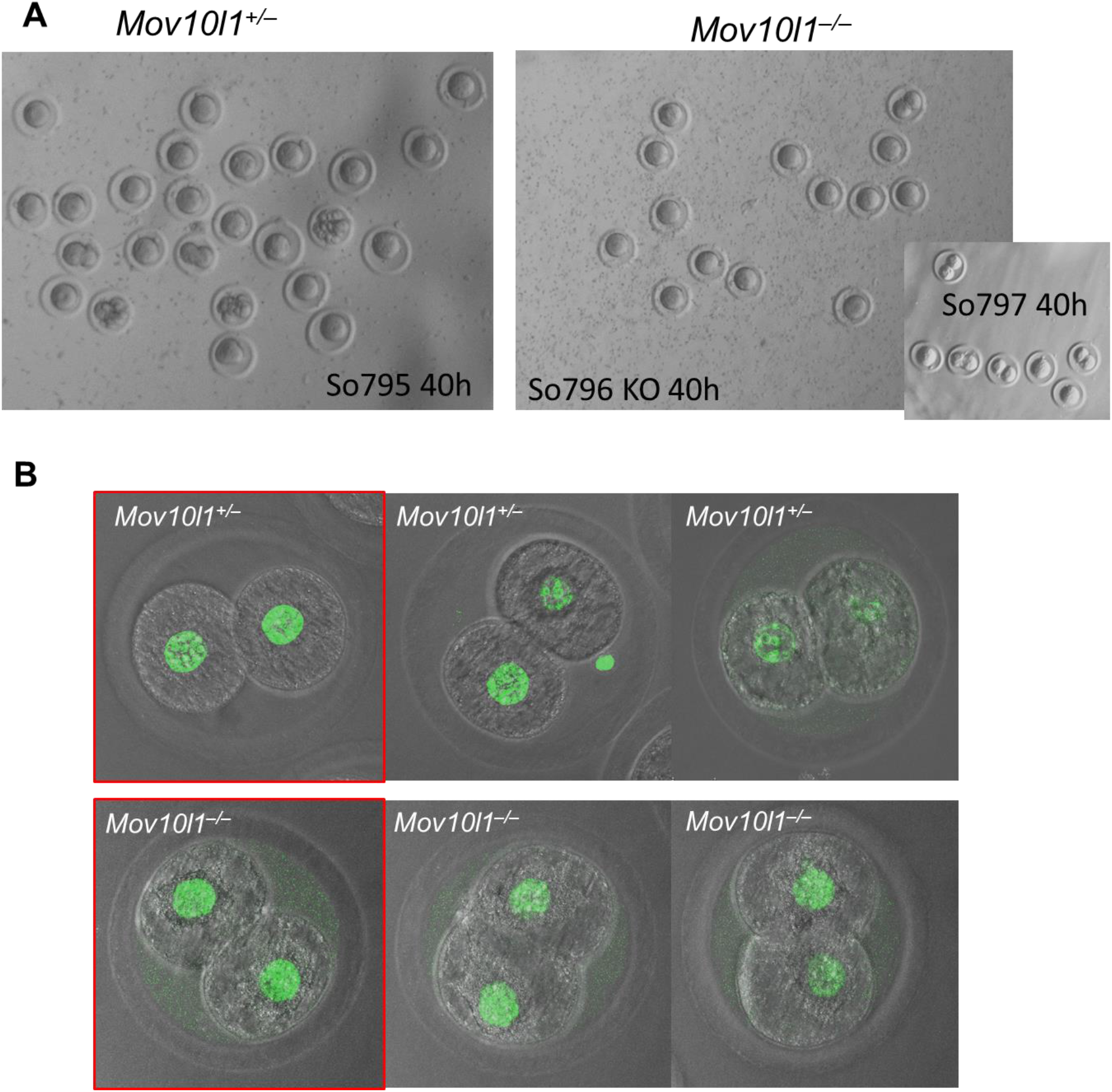
*Mov10l1^−/−^* oocytes can give rise to 2-cell zygotes. (A) View of fertilized eggs and cleaving embryos isolated from *Mov10l1^+/−^* and *Mov10l1^−/−^* females 40 h after mating. (B) Various 2-cell zygotes. Green nuclear signal is H3K9me3 staining, which shows no difference between *Mov10l1^+/-^* and *Mov10l1^−/−^* 2-cell zygotes.

**Figure S8.**
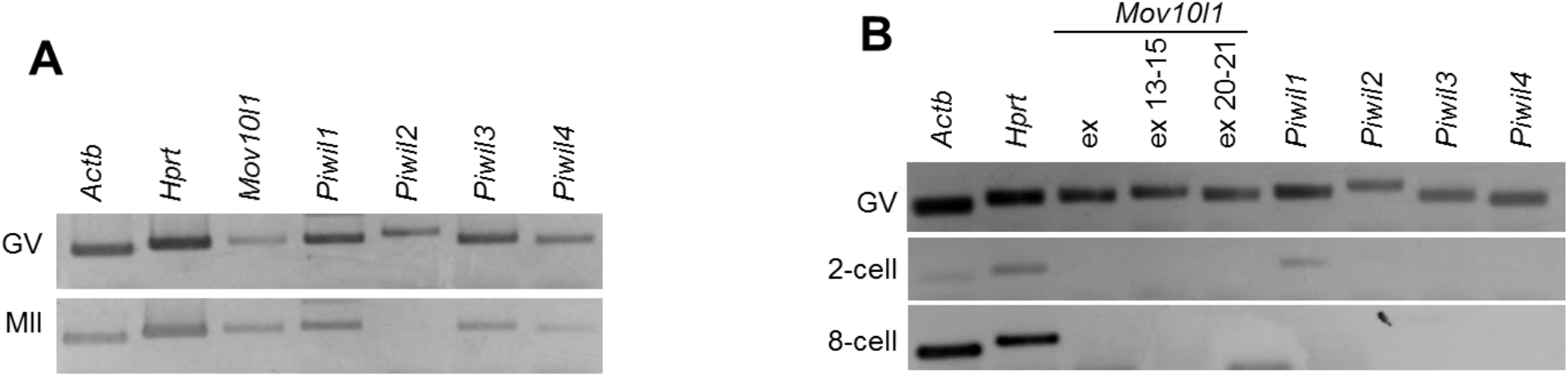
Expression patterns of genes encoding *MOV10L1* and *PIWI* proteins suggest that mRNAs encoding *MOV10L1* and *PIWI* proteins become degraded during meiotic maturation (A) and degradation of these maternal mRNAs is mostly finished by the 2-cells stage. (B) Equal cDNA “embryo-equivalent” amount was used for each PCR. Thus, reduced amplification of *Actb* and *Hprt* in MII and 2-cell stages presumably reflects maternal mRNA degradation of these housekeeping transcripts.

**Figure S9.**
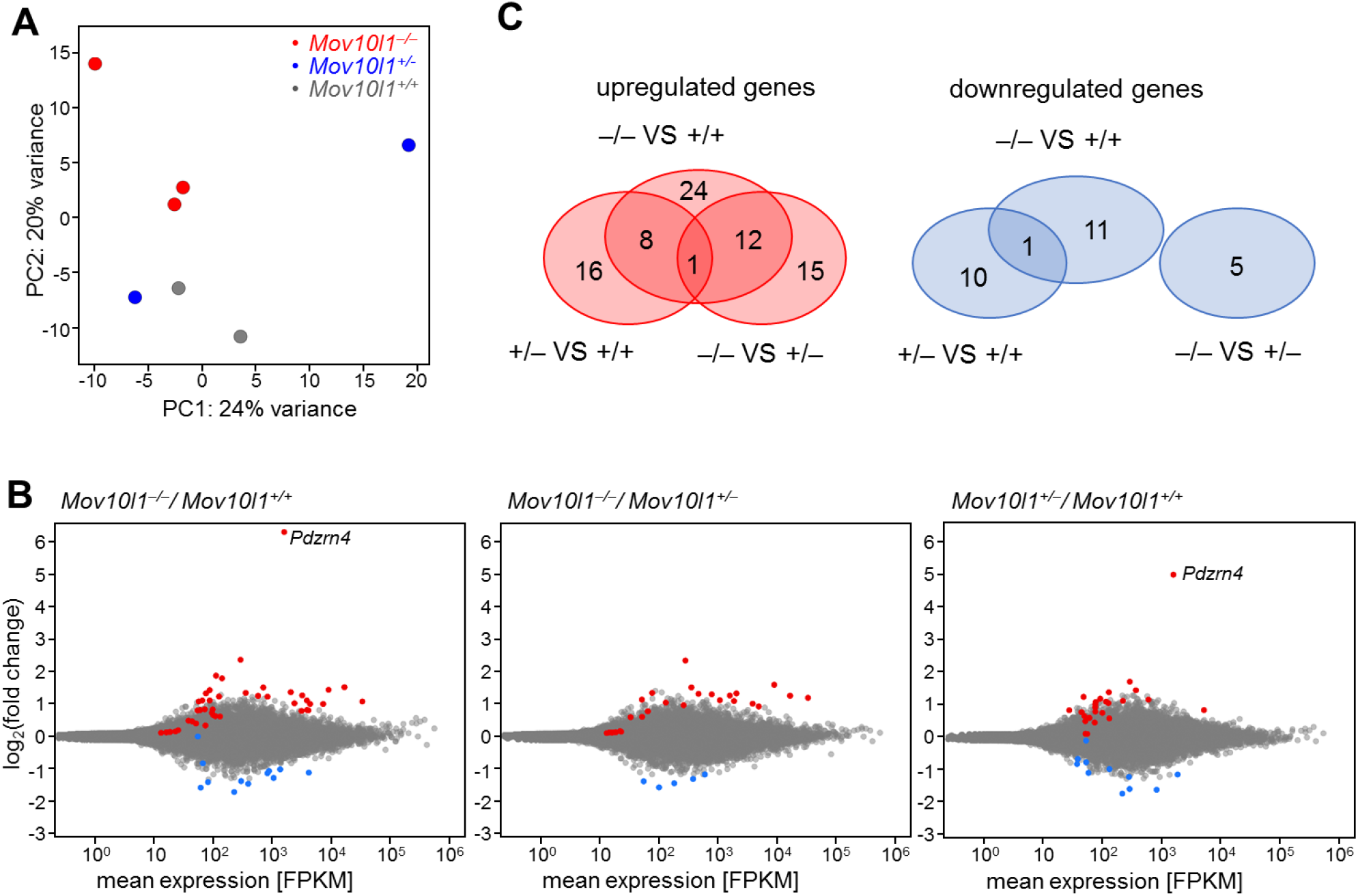
Transcriptome changes in *Mov10l1^−/−^* oocytes. (A) PCA analysis depicting distribution of individual sequencing libraries. (B) MA plots for all genotype combinations. The MA plot on the left is shown as Fig. 3E. Because of the trimming, *Pdzrn4* is not displayed in Fig. 3E. However, it’s upregulation is not specific to *Mov10l1^−/−^* oocytes because it is high also in *Mov10l^+/−^* oocytes. Red and blue points depict protein-coding genes with significantly higher and lower transcript abundance in *Mov10l1^−/−^* oocytes, respectively (DESeq2 p-value <0.01). (C) Venn diagrams show overlaps among transcripts with increased or decreased relative abundance.

**Figure S10.**
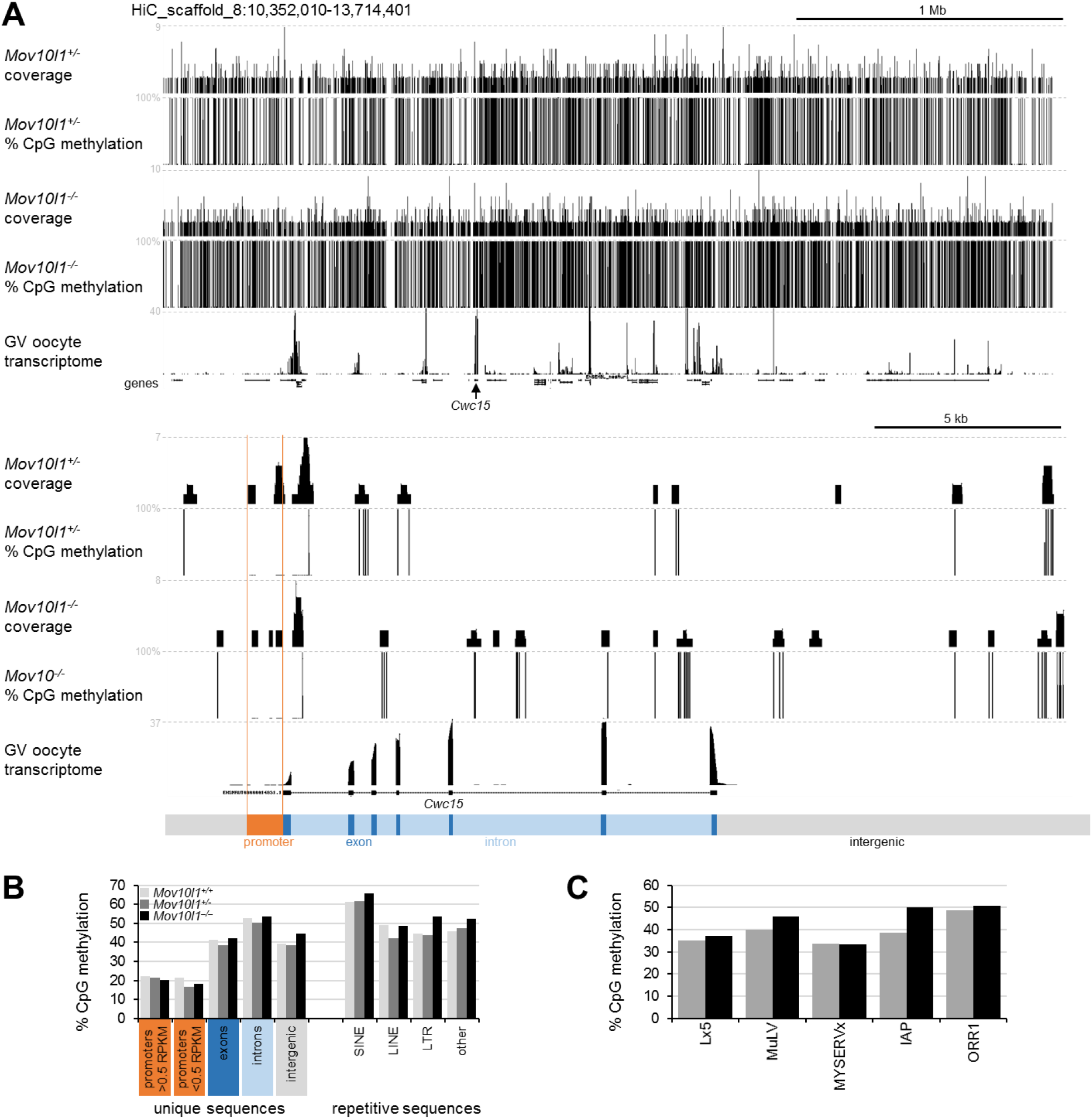
Bisulfite sequencing analysis of *Mov10l1* mutants. (A) Snapshots from the UCSC genome browser depicting coverage of the genome by bisulfite-sequenced fragments and CpG methylation frequency. The upper panel depicts a larger genomic region, the lower panel depicts expressed Cwc15 gene with apparent absence of DNA methylation in the promoter. (B) Quantification of distribution of CpG methylation in hamster oocytes in unique and selected repetitive sequences. Only regions covered by at least four fragments in *Mov10l1^+/−^* and *Mov10l1^−/−^* libraries were included in the analysis. Promoters were considered regions 1 kb upstream of an annotated transcription start sites and were divided into two groups according to their activity using an arbitrary expression threshold of 0.5 RPKM. (C) CpG methylation of selected retrotransposon subgroups.

**Figure S11.**
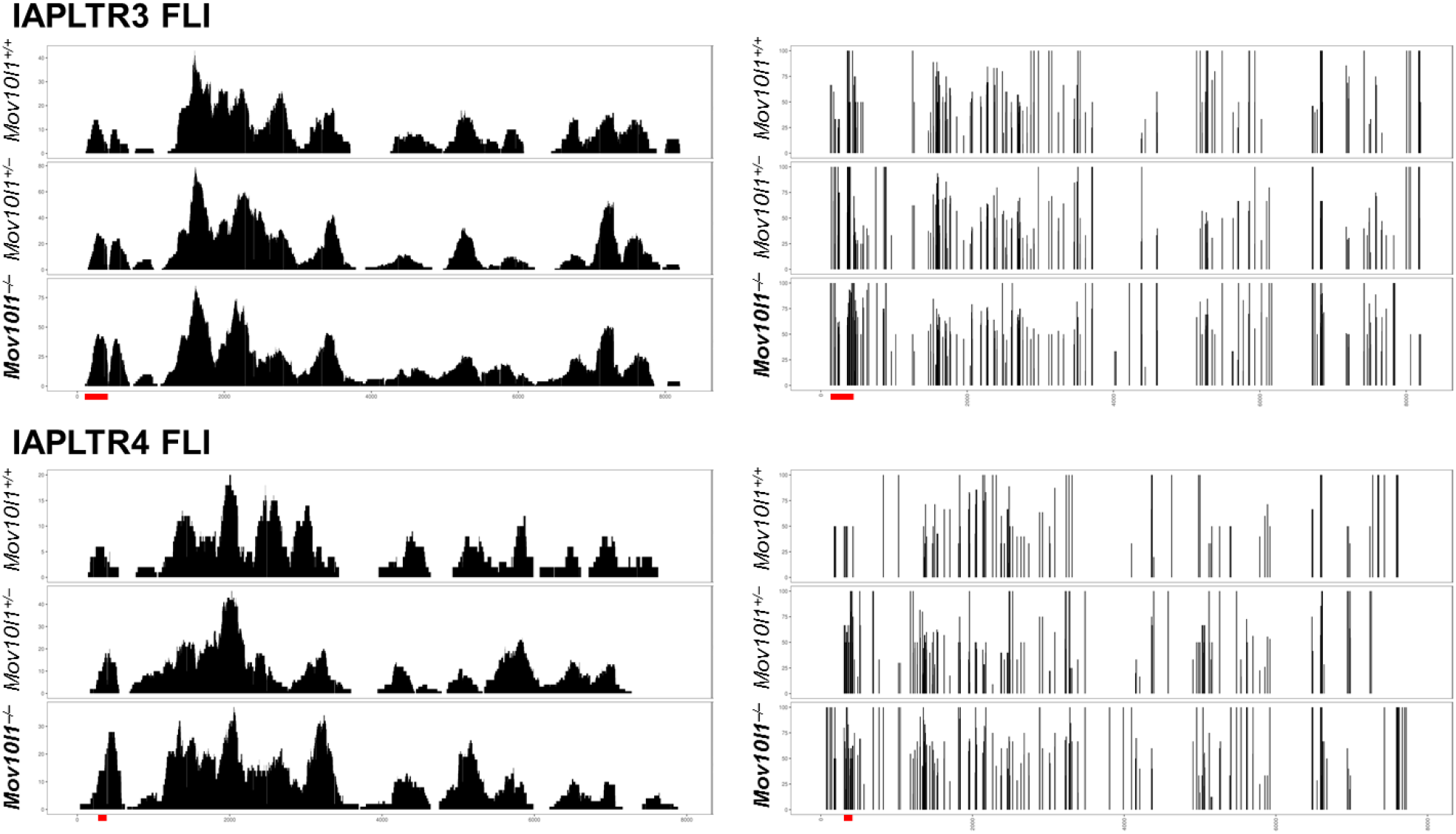
CpG methylation in IAP full-length intact retrotransposons. Panels on the left depict “coverage” of the retrotransposon genome, panels on the right display methylation frequency at CpGs (vertical lines). Different coverage of 5’ and 3’ LTR sequences (which are identical) comes from the fact that sequencing was done as “paired-end sequencing”, hence if one read mapped exclusively to an LTR, the second read typically allowed to distinguish between 3’ and 5’ LTRs. Red lines depict CpG positions displayed in Fig. 3H.

**Figure S12.**
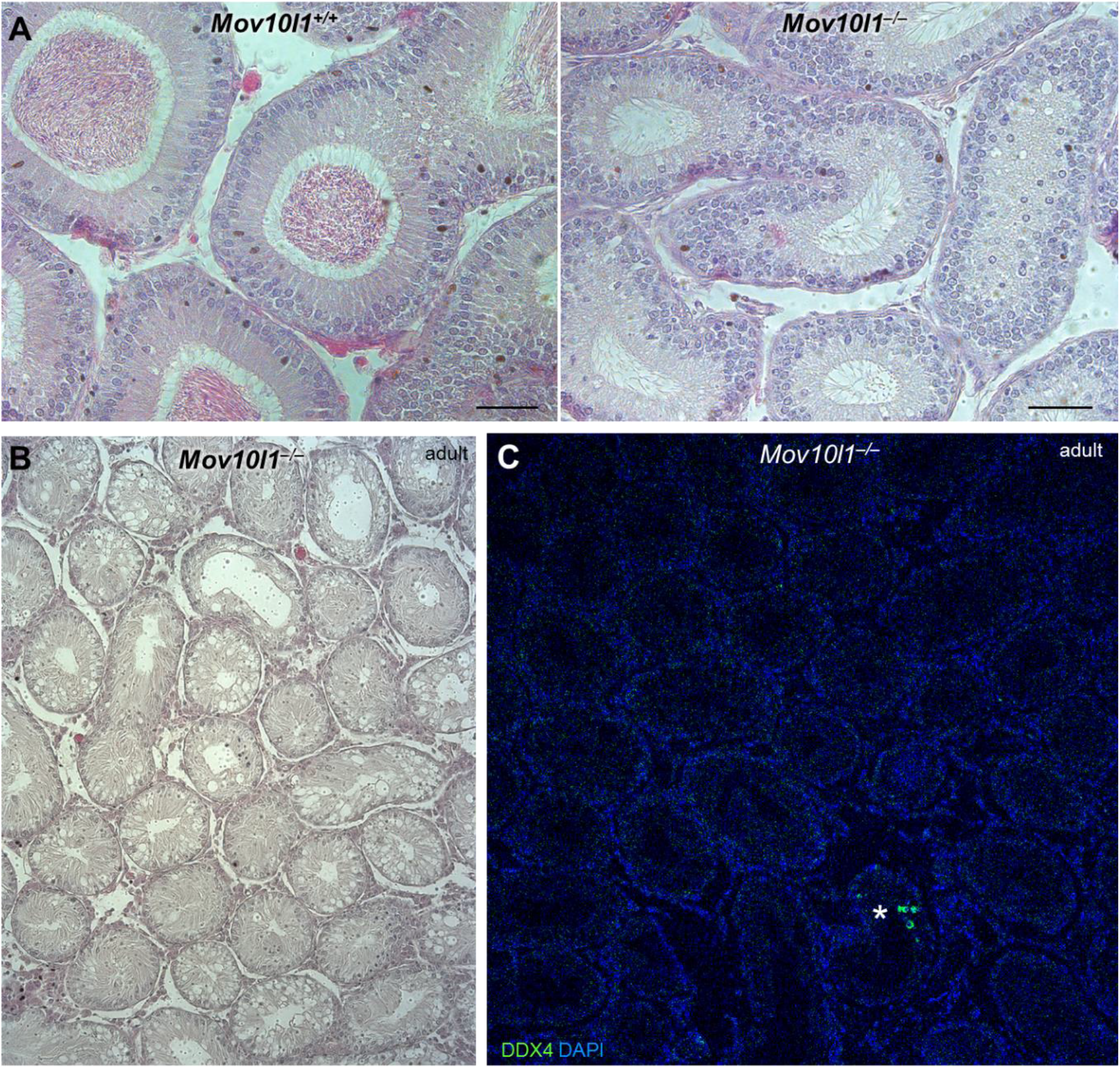
Sterility in *Mov10l1^−/−^* males. (A) H&E staining of epididymal sections shows lack of sperm production in adult *Mov10l1^−/−^* testes. Scale bar = 50 μm. (B) Most of the testicular seminiferous tubules in *Mov10l1^−/−^* testes appear aspermatogenic. (C) Immunofluorescent staining with germ cell marker DDX4 (VASA) reveals rare clusters of spermatogenic cells (97% of tubules do not contain any germ cells).

**Figure S13.**
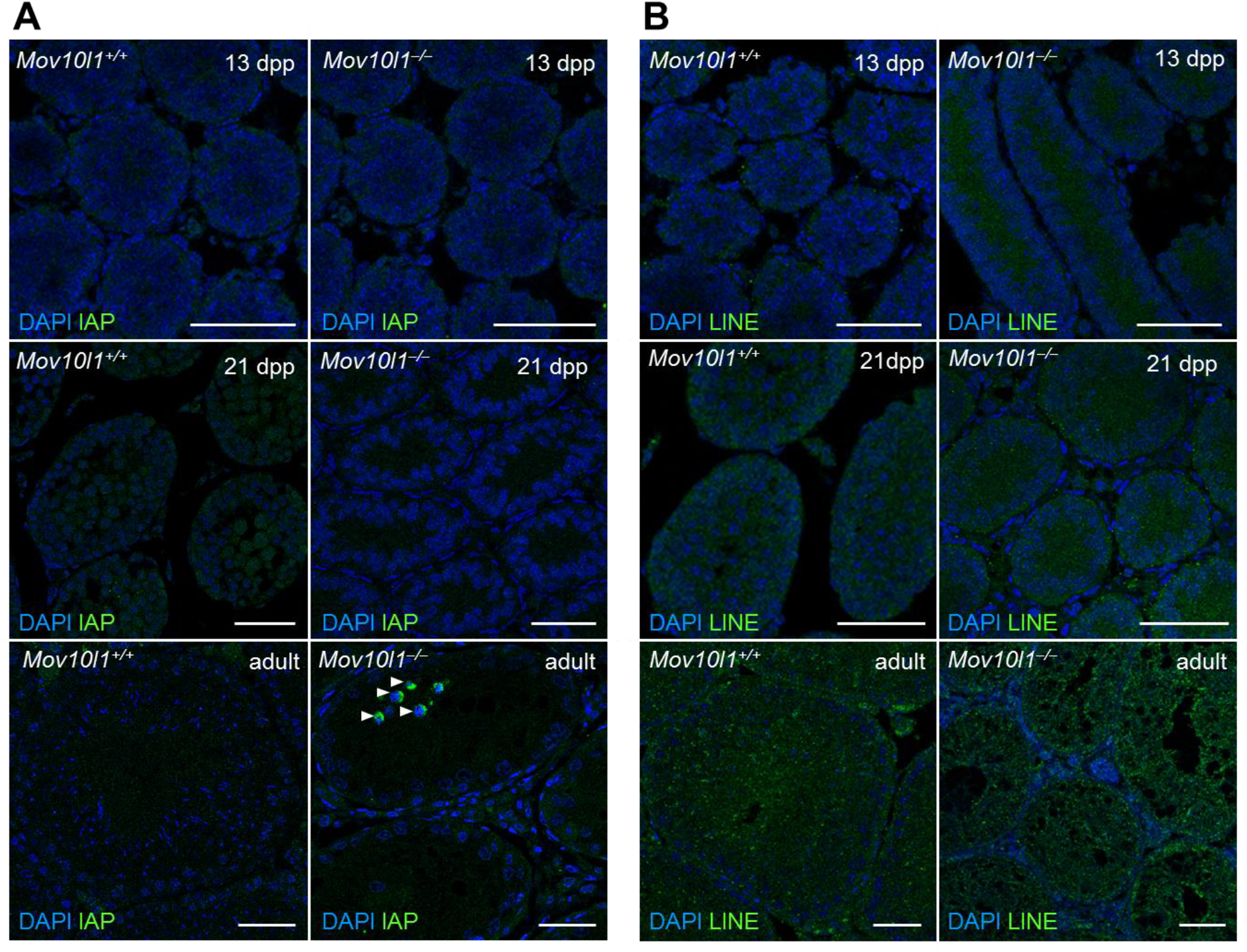
Analysis of LINE1 and IAP expression in 13 dpp, 21 dpp, and adult testes did not reveal any retrotransposon mobilization upon loss of *Mov10l1^−/−^*. (A) IAP showed only background signal at 13 dpp and 21 dpp, but positive signal in the aforementioned clusters of spermatogenic cells in adults (pointed at by arrowheads). (B) LINE1 showed only background signal at 13 dpp, 21 dpp and adults, but we cannot rule out that the antibody against ORF1 of mouse LINE1 did not react with Syrian hamster LINE elements. Scale bars = 50 μm.

**Figure S14.**
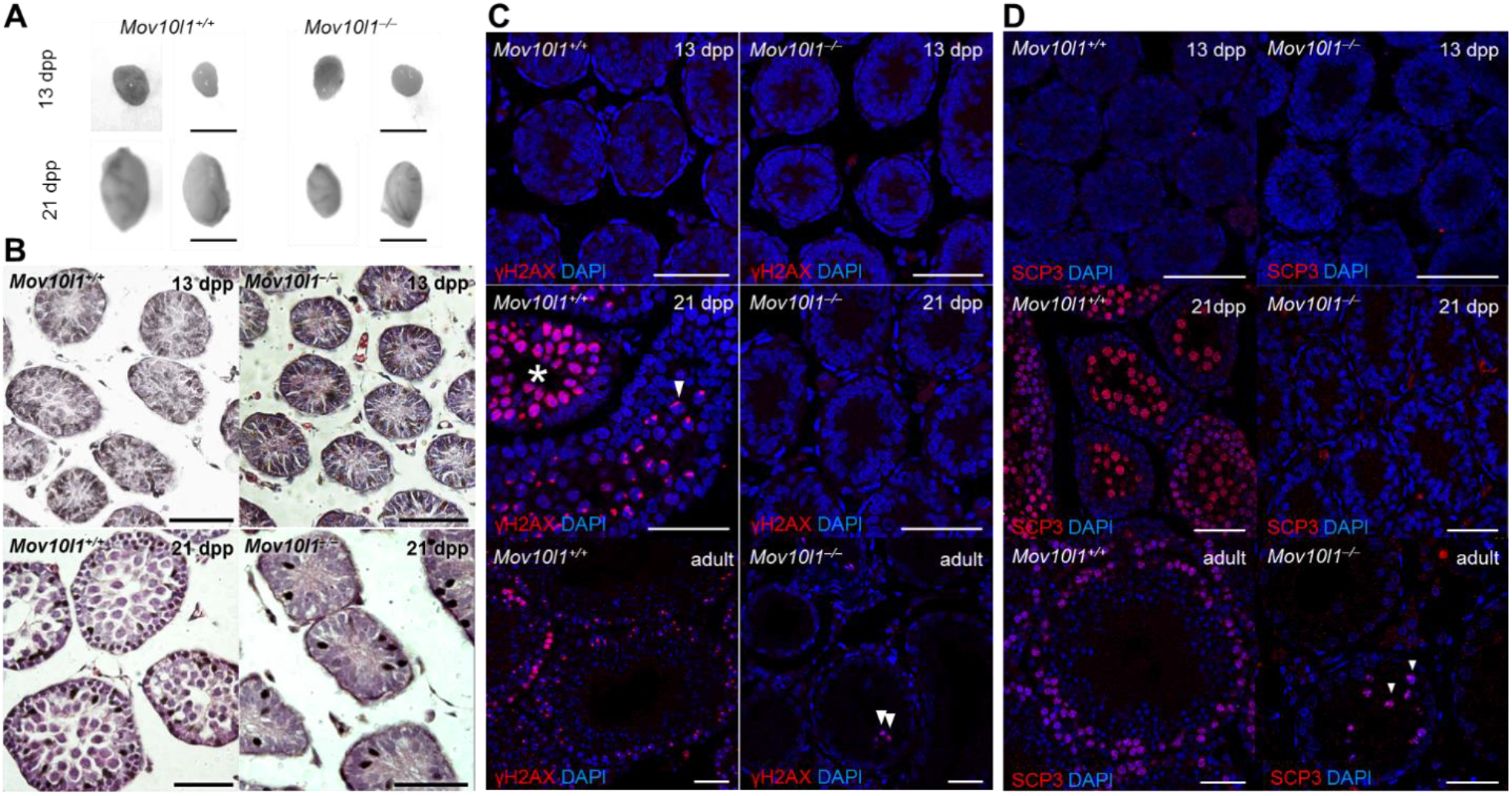
Spermatogenesis defects in *Mov10l1^−/−^* testes. (A) 21 dpp *Mov10l1^−/−^* testes were visibly smaller than normal testes. (B) Haematoxylin-eosin staining reveals major defects in the seminiferous tubule appearance in mutants. (C) γH2AX staining suggests absence of meiotic cells at 21 dpp and adult testes, 13 dpp testes were used as a negative control. (D) 21 dpp testes lack entirely meiotic cells as shown by SCP3 staining, which is a marker of meiotic cells. Adult testes show rare positive staining in the aforementioned clusters (Fig. 3C), 13 dpp testes were used as a negative control. Scale bars = 50 μm.

**Figure S15.**
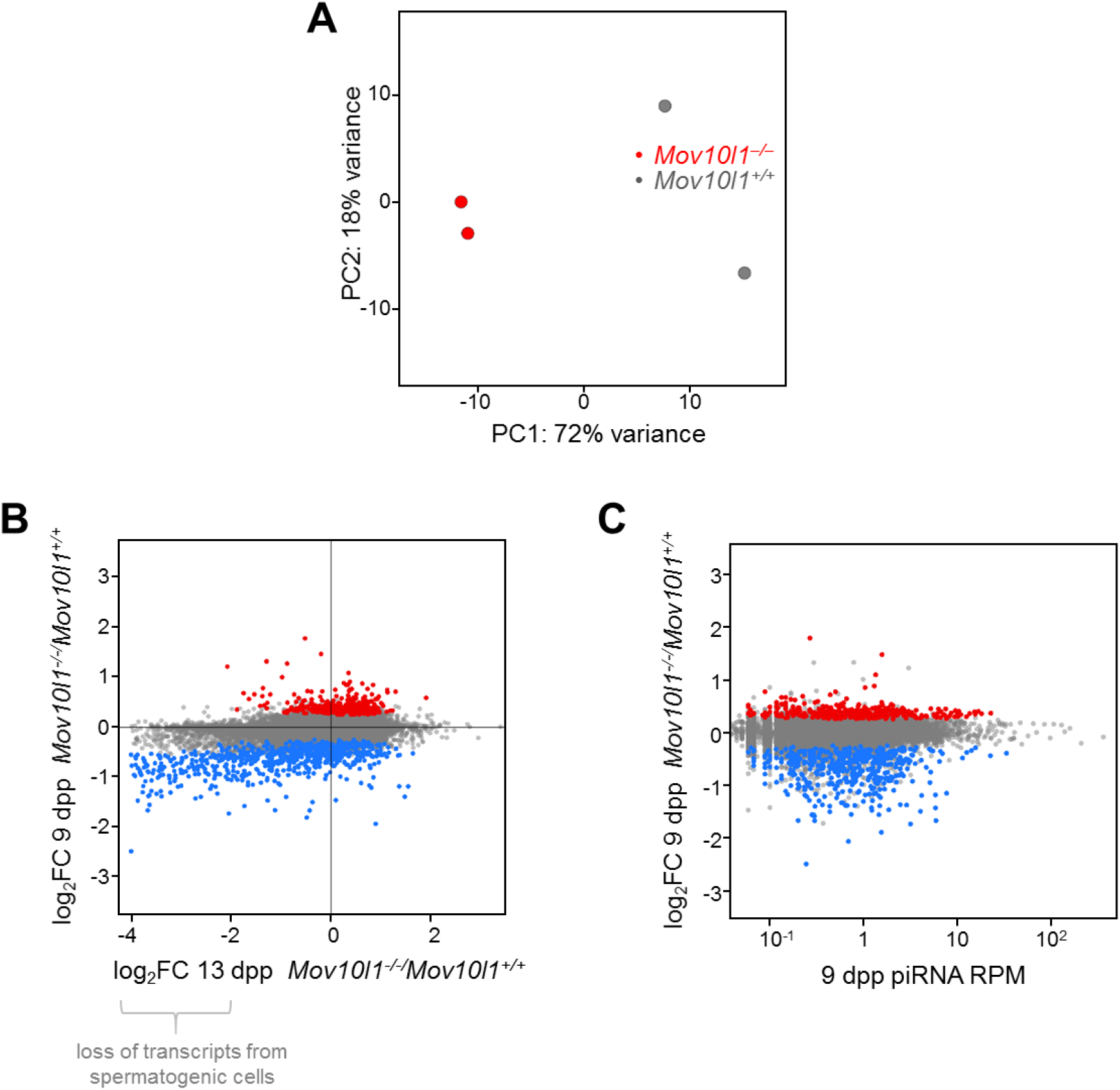
Transcriptome changes in *Mov10l1^−/−^* 9 dpp testes. (A) PCA analysis depicting distribution of individual sequencing libraries. (B) Relative gene expression changes in *Mov10l1^−/−^* 9 dpp testes (y-axis) plotted against relative difference of *Mov10l1^+/+^* and *Mov10l1^−/−^* 13 dpp testes (x-axis). Since 13 dpp *Mov10l1^−/−^* testes lack most spermatogonia, comparison of normal and knock-out 13 dpp testes reveals loss of expression of genes whose expression was restricted to germ cells. Thus, points (significantly downregulated genes in *Mov10l1^−/−^* 9 dpp testes) that localize to the left side of the plot, visualize germ cell-specific genes are disturbed at 9 dpp. (C) Relative gene expression changes in *Mov10l1^−/−^* 9 dpp testes (y-axis) plotted against rank-sorted abundance of 24-31 nt reads mapping to the transcribed regions of these genes. A fraction of differentially expressed genes associated with piRNAs over 5 RPM are typically upregulated.

**Figure S16.**
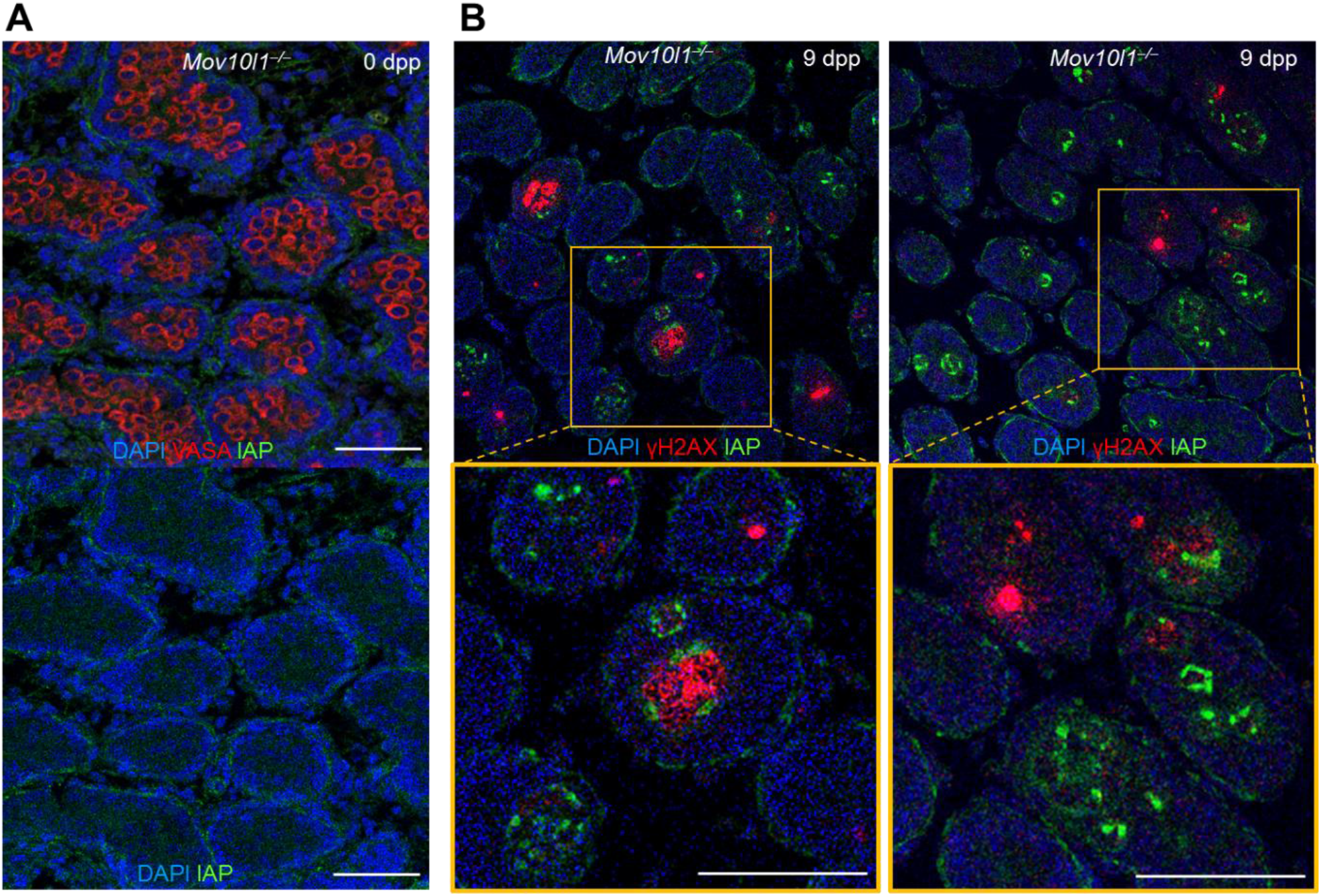
Increased expression of IAP GAG protein expression in 9 dpp testes. (A) 0 dpp *Mov10l1^−/−^* testes shows normal presence of the germ cell marker DDX4 (VASA) and no mobilization of IAP expression in germ cells. (B) Fluorescence staining of IAP GAG and γH2AX reveals that increased IAP expression in germ cells in seminiferous tubules is not accompanied by immediate formation of γH2AX foci in the nucleus. Scale bars = 50 μm.

**Table S1** 9 dpp pre-pachytene piRNA clusters

**Table S2** 13 dpp pre-pachytene piRNA clusters

**Table S3** 21 dpp pachytene piRNA clusters

**Table S4** DEG genes in *Mov10l1^−/−^* GV oocytes

**Table S5** DEG genes in *Mov10l1^−/−^* 9 dpp testes

**Table S6.**
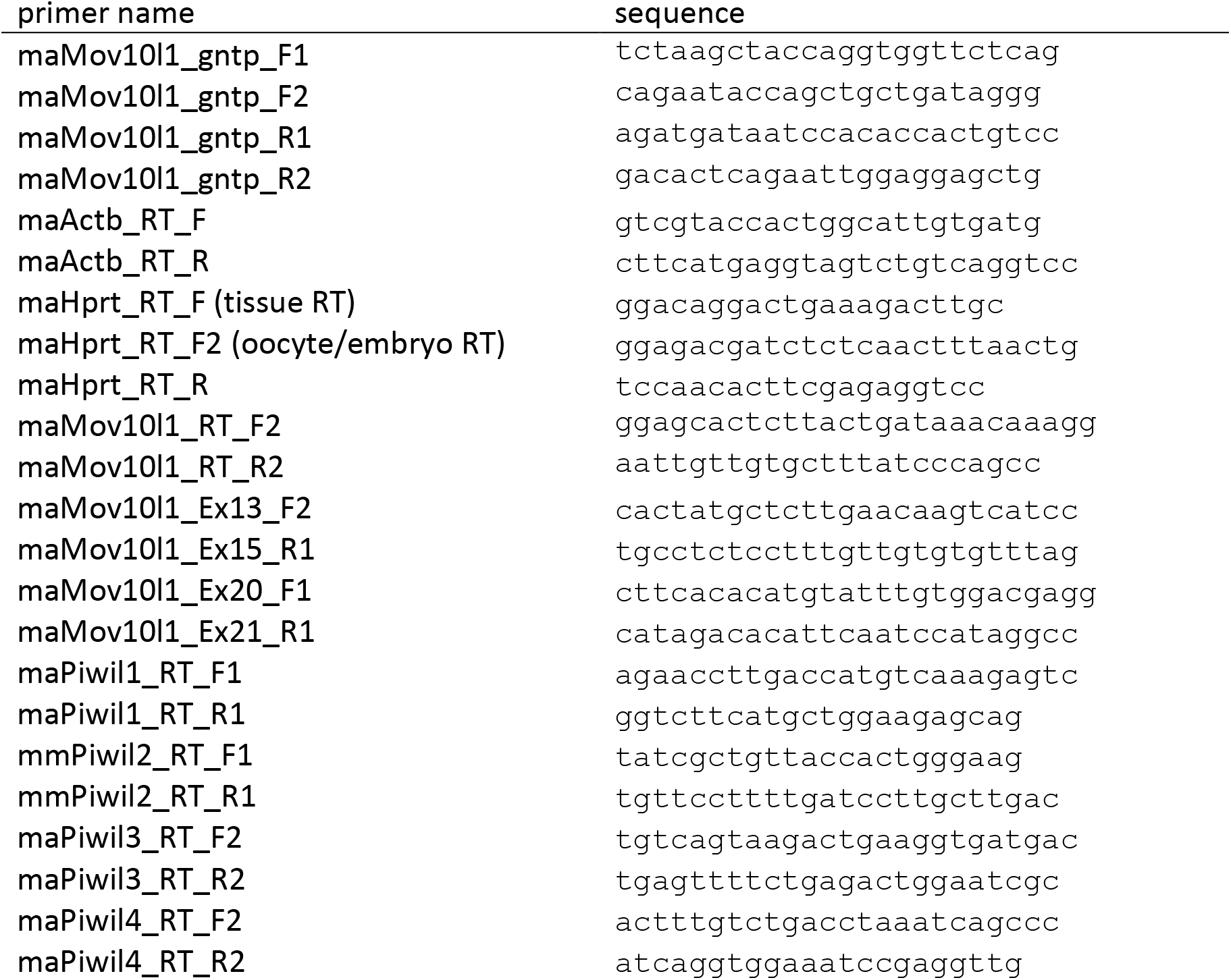
Primers

**Table S7.**
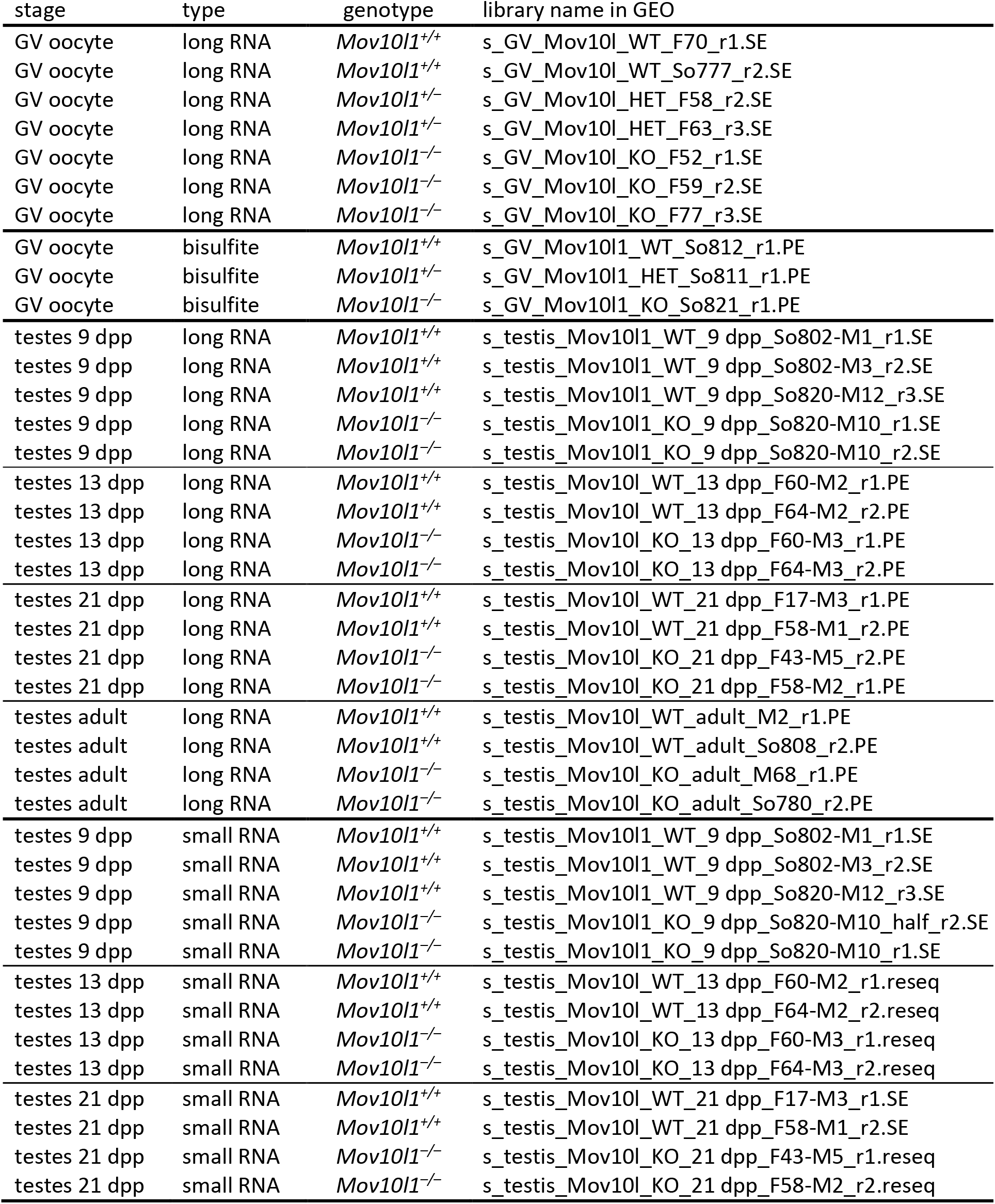
List of RNA-seq libraries used in this study

| stage | type | genotype | library name in GEO |
| --- | --- | --- | --- |
| GV oocyte | long RNA | *Mov10l1*<sup>+/+</sup> | s_GV_Mov10l_WT_F70_r1.SE |
| GV oocyte | long RNA | *Mov10l1*<sup>+/+</sup> | s_GV_Mov10l_WT_So777_r2.SE |
| GV oocyte | long RNA | *Mov10l1*<sup>+/-</sup> | s_GV_Mov10l_HET_F58_r2.SE |
| GV oocyte | long RNA | *Mov10l1*<sup>+/-</sup> | s_GV_Mov10l_HET_F63_r3.SE |
| GV oocyte | long RNA | *Mov10l1*<sup>-/-</sup> | s_GV_Mov10l_KO_F52_r1.SE |
| GV oocyte | long RNA | *Mov10l1*<sup>-/-</sup> | s_GV_Mov10l_KO_F59_r2.SE |
| GV oocyte | long RNA | *Mov10l1*<sup>-/-</sup> | s_GV_Mov10l_KO_F77_r3.SE |
| GV oocyte | bisulfite | *Mov10l1*<sup>+/+</sup> | s_GV_Mov10l1_WT_So812_r1.PE |
| GV oocyte | bisulfite | *Mov10l1*<sup>+/-</sup> | s_GV_Mov10l1_HET_So811_r1.PE |
| GV oocyte | bisulfite | *Mov10l1*<sup>-/-</sup> | s_GV_Mov10l1_KO_So821_r1.PE |
| testes 9 dpp | long RNA | *Mov10l1*<sup>+/+</sup> | s_testis_Mov10l1_WT_9 dpp_So802-M1_r1.SE |
| testes 9 dpp | long RNA | *Mov10l1*<sup>+/+</sup> | s_testis_Mov10l1_WT_9 dpp_So802-M3_r2.SE |
| testes 9 dpp | long RNA | *Mov10l1*<sup>+/+</sup> | s_testis_Mov10l1_WT_9 dpp_So820-M12_r3.SE |
| testes 9 dpp | long RNA | *Mov10l1*<sup>-/-</sup> | s_testis_Mov10l1_KO_9 dpp_So820-M10_r1.SE |
| testes 9 dpp | long RNA | *Mov10l1*<sup>-/-</sup> | s_testis_Mov10l1_KO_9 dpp_So820-M10_r2.SE |
| testes 13 dpp | long RNA | *Mov10l1*<sup>+/+</sup> | s_testis_Mov10l_WT_13 dpp_F60-M2_r1.PE |
| testes 13 dpp | long RNA | *Mov10l1*<sup>+/+</sup> | s_testis_Mov10l_WT_13 dpp_F64-M2_r2.PE |
| testes 13 dpp | long RNA | *Mov10l1*<sup>-/-</sup> | s_testis_Mov10l_KO_13 dpp_F60-M3_r1.PE |
| testes 13 dpp | long RNA | *Mov10l1*<sup>-/-</sup> | s_testis_Mov10l_KO_13 dpp_F64-M3_r2.PE |
| testes 21 dpp | long RNA | *Mov10l1*<sup>+/+</sup> | s_testis_Mov10l_WT_21 dpp_F17-M3_r1.PE |
| testes 21 dpp | long RNA | *Mov10l1*<sup>+/+</sup> | s_testis_Mov10l_WT_21 dpp_F58-M1_r2.PE |
| testes 21 dpp | long RNA | *Mov10l1*<sup>-/-</sup> | s_testis_Mov10l_KO_21 dpp_F43-M5_r2.PE |
| testes 21 dpp | long RNA | *Mov10l1*<sup>-/-</sup> | s_testis_Mov10l_KO_21 dpp_F58-M2_r1.PE |
| testes adult | long RNA | *Mov10l1*<sup>+/+</sup> | s_testis_Mov10l_WT_adult_M2_r1.PE |
| testes adult | long RNA | *Mov10l1*<sup>+/+</sup> | s_testis_Mov10l_WT_adult_So808_r2.PE |
| testes adult | long RNA | *Mov10l1*<sup>-/-</sup> | s_testis_Mov10l_KO_adult_M68_r1.PE |
| testes adult | long RNA | *Mov10l1*<sup>-/-</sup> | s_testis_Mov10l_KO_adult_So780_r2.PE |
| testes 9 dpp | small RNA | *Mov10l1*<sup>+/+</sup> | s_testis_Mov10l1_WT_9 dpp_So802-M1_r1.SE |
| testes 9 dpp | small RNA | *Mov10l1*<sup>+/+</sup> | s_testis_Mov10l1_WT_9 dpp_So802-M3_r2.SE |
| testes 9 dpp | small RNA | *Mov10l1*<sup>+/+</sup> | s_testis_Mov10l1_WT_9 dpp_So820-M12_r3.SE |
| testes 9 dpp | small RNA | *Mov10l1*<sup>-/-</sup> | s_testis_Mov10l1_KO_9 dpp_So820-M10_half_r2.SE |
| testes 9 dpp | small RNA | *Mov10l1*<sup>-/-</sup> | s_testis_Mov10l1_KO_9 dpp_So820-M10_r1.SE |
| testes 13 dpp | small RNA | *Mov10l1*<sup>+/+</sup> | s_testis_Mov10l_WT_13 dpp_F60-M2_r1.reseq |
| testes 13 dpp | small RNA | *Mov10l1*<sup>+/+</sup> | s_testis_Mov10l_WT_13 dpp_F64-M2_r2.reseq |
| testes 13 dpp | small RNA | *Mov10l1*<sup>-/-</sup> | s_testis_Mov10l_KO_13 dpp_F60-M3_r1.reseq |
| testes 13 dpp | small RNA | *Mov10l1*<sup>-/-</sup> | s_testis_Mov10l_KO_13 dpp_F64-M3_r2.reseq |
| testes 21 dpp | small RNA | *Mov10l1*<sup>+/+</sup> | s_testis_Mov10l_WT_21 dpp_F17-M3_r1.SE |
| testes 21 dpp | small RNA | *Mov10l1*<sup>+/+</sup> | s_testis_Mov10l_WT_21 dpp_F58-M1_r2.SE |
| testes 21 dpp | small RNA | *Mov10l1*<sup>-/-</sup> | s_testis_Mov10l_KO_21 dpp_F43-M5_r1.reseq |
| testes 21 dpp | small RNA | *Mov10l1*<sup>-/-</sup> | s_testis_Mov10l_KO_21 dpp_F58-M2_r2.reseq |

**Data S1** ERVK LTR retrotransposon nucleotide exchange rates

**Data S2** Other LTR retrotransposon nucleotide exchange rates

**Data S3** MaLR LTR retrotransposon nucleotide exchange rates

**Data S4** Golden hamster IAP FLI sequences

**Data S5** Golden hamster LINE L1 FLI sequences

## REFERENCES

1 Aravin, A.A., Hannon, G.J. and Brennecke, J. (2007) The Piwi-piRNA pathway provides an adaptive defense in the transposon arms race. Science, 318, 761–764.

2 Ozata, D.M., Gainetdinov, I., Zoch, A., O’Carroll, D. and Zamore, P.D. (2019) PIWI-interacting RNAs: small RNAs with big functions. Nat Rev Genet, 20, 89–108.

3 Frost, R.J., Hamra, F.K., Richardson, J.A., Qi, X., Bassel-Duby, R. and Olson, E.N. (2010) MOV10L1 is necessary for protection of spermatocytes against retrotransposons by Piwi-interacting RNAs. Proc Natl Acad Sci U S A, 107, 11847–11852.

4 Zheng, K., Xiol, J., Reuter, M., Eckardt, S., Leu, N.A., McLaughlin, K.J., Stark, A., Sachidanandam, R., Pillai, R.S. and Wang, P.J. (2010) Mouse MOV10L1 associates with Piwi proteins and is an essential component of the Piwi-interacting RNA (piRNA) pathway. Proceedings of the National Academy of Sciences of the United States of America, 107, 11841–11846.

5 Vourekas, A., Zheng, K., Fu, Q., Maragkakis, M., Alexiou, P., Ma, J., Pillai, R.S., Mourelatos, Z. and Wang, P.J. (2015) The RNA helicase MOV10L1 binds piRNA precursors to initiate piRNA processing. Genes & Development, 29, 617–629.

6 Aravin, A.A., Sachidanandam, R., Girard, A., Fejes-Toth, K. and Hannon, G.J. (2007) Developmentally regulated piRNA clusters implicate MILI in transposon control. Science, 316, 744–747.

7 Kuramochi-Miyagawa, S., Watanabe, T., Gotoh, K., Totoki, Y., Toyoda, A., Ikawa, M., Asada, N., Kojima, K., Yamaguchi, Y., Ijiri, T.W. et al. (2008) DNA methylation of retrotransposon genes is regulated by Piwi family members MILI and MIWI2 in murine fetal testes. Genes Dev, 22, 908–917.

8 Robine, N., Lau, N.C., Balla, S., Jin, Z., Okamura, K., Kuramochi-Miyagawa, S., Blower, M.D. and Lai, E.C. (2009) A broadly conserved pathway generates 3’UTR-directed primary piRNAs. Curr Biol, 19, 2066–2076.

9 Chirn, G.W., Rahman, R., Sytnikova, Y.A., Matts, J.A., Zeng, M., Gerlach, D., Yu, M., Berger, B., Naramura, M., Kile, B.T. et al. (2015) Conserved piRNA Expression from a Distinct Set of piRNA Cluster Loci in Eutherian Mammals. PLoS Genet, 11, e1005652.

10 Yang, Q., Li, R., Lyu, Q., Hou, L., Liu, Z., Sun, Q., Liu, M., Shi, H., Xu, B., Yin, M. et al. (2019) Single-cell CAS-seq reveals a class of short PIWI-interacting RNAs in human oocytes. Nature communications, 10, 3389.

11 Meister, G. (2013) Argonaute proteins: functional insights and emerging roles. Nat Rev Genet, 14, 447–459.

12 Kuramochi-Miyagawa, S., Kimura, T., Ijiri, T.W., Isobe, T., Asada, N., Fujita, Y., Ikawa, M., Iwai, N., Okabe, M., Deng, W. et al. (2004) Mili, a mammalian member of piwi family gene, is essential for spermatogenesis. Development, 131, 839–849.

13 Deng, W. and Lin, H.F. (2002) miwi, a murine homolog of piwi, encodes a cytoplasmic protein essential for spermatogenesis. Developmental Cell, 2, 819–830.

14 Carmell, M.A., Girard, A., van de Kant, H.J.G., Bourc’his, D., Bestor, T.H., de Rooij, D.G. and Hannon, G.J. (2007) MIWI2 is essential for spermatogenesis and repression of transposons in the mouse male germline. Developmental Cell, 12, 503–514.

15 Lim, A.K., Lorthongpanich, C., Chew, T.G., Tan, C.W., Shue, Y.T., Balu, S., Gounko, N., Kuramochi-Miyagawa, S., Matzuk, M.M., Chuma, S. et al. (2013) The nuage mediates retrotransposon silencing in mouse primordial ovarian follicles. Development, 140, 3819–3825.

16 Kabayama, Y., Toh, H., Katanaya, A., Sakurai, T., Chuma, S., Kuramochi-Miyagawa, S., Saga, Y., Nakano, T. and Sasaki, H. (2017) Roles of MIWI, MILI and PLD6 in small RNA regulation in mouse growing oocytes. Nucleic Acids Research, 45, 5387–5398.

17 Flemr, M., Malik, R., Franke, V., Nejepinska, J., Sedlacek, R., Vlahovicek, K. and Svoboda, P. (2013) A Retrotransposon-Driven Dicer Isoform Directs Endogenous Small Interfering RNA Production in Mouse Oocytes. Cell, 155, 807–816.

18 Taborska, E., Pasulka, J., Malik, R., Horvat, F., Jenickova, I., Jelic Matosevic, Z. and Svoboda, P. (2019) Restricted and non-essential redundancy of RNAi and piRNA pathways in mouse oocytes. PLoS Genet, 15, e1008261.

19 Roovers, E.F., Rosenkranz, D., Mahdipour, M., Han, C.T., He, N., Chuva de Sousa Lopes, S.M., van der Westerlaken, L.A., Zischler, H., Butter, F., Roelen, B.A. et al. (2015) Piwi proteins and piRNAs in mammalian oocytes and early embryos. Cell Rep, 10, 2069–2082.

20 Steppan, S., Adkins, R. and Anderson, J. (2004) Phylogeny and divergence-date estimates of rapid radiations in muroid rodents based on multiple nuclear genes. Syst Biol, 53, 533–553.

21 Hirose, M. and Ogura, A. (2019) The golden (Syrian) hamster as a model for the study of reproductive biology: Past, present, and future. Reprod Med Biol, 18, 34–39.

22 Franke, V., Ganesh, S., Karlic, R., Malik, R., Pasulka, J., Horvat, F., Kuzman, M., Fulka, H., Cernohorska, M., Urbanova, J. et al. (2017) Long terminal repeats power evolution of genes and gene expression programs in mammalian oocytes and zygotes. Genome Research, 27, 1384–1394.

23 Hirose, M., Honda, A., Fulka, H., Tamura-Nakano, M., Matoba, S., Tomishima, T., Mochida, K., Hasegawa, A., Nagashima, K., Inoue, K. et al. (2020) Acrosin is essential for sperm penetration through the zona pellucida in hamsters. Proc Natl Acad Sci U S A, 117, 2513–2518.

24 Ishino, K., Hasuwa, H., Yoshimura, J., Iwasaki, Y.W., Nishihara, H., Seki, N.M., Hirano, T., Tsuchiya, M., Ishizaki, H., Masuda, H. et al. (2020) Hamster PIWI proteins bind to piRNAs with stage-specific size variations during oocyte maturation. bioRxiv, 2020.2012.2001.407411.

25 Miething, A. (1998) The establishment of spermatogenesis in the seminiferous epithelium of the pubertal golden hamster (Mesocricetus auratus). Adv Anat Embryol Cell Biol, 140, 1–92.

26 Ribet, D., Harper, F., Dupressoir, A., Dewannieux, M., Pierron, G. and Heidmann, T. (2008) An infectious progenitor for the murine IAP retrotransposon: emergence of an intracellular genetic parasite from an ancient retrovirus. Genome Res, 18, 597–609.

27 Magiorkinis, G., Gifford, R.J., Katzourakis, A., De Ranter, J. and Belshaw, R. (2012) Env-less endogenous retroviruses are genomic superspreaders. Proc Natl Acad Sci U S A, 109, 7385–7390.

28 Dewannieux, M., Dupressoir, A., Harper, F., Pierron, G. and Heidmann, T. (2004) Identification of autonomous IAP LTR retrotransposons mobile in mammalian cells. Nat Genet, 36, 534–539.

29 Ostertag, E.M. and Kazazian, H.H. Jr. (2001) Biology of mammalian L1 retrotransposons. Annu Rev Genet, 35, 501–538.

30 Penzkofer, T., Jager, M., Figlerowicz, M., Badge, R., Mundlos, S., Robinson, P.N. and Zemojtel, T. (2017) L1Base 2: more retrotransposition-active LINE-1s, more mammalian genomes. Nucleic Acids Res, 45, D68–D73.

31 Hasuwa, H. Y.W. I., Wan Kin, A.Y., Ishino, K., Masuda, H., Sasaki, H. and Siomi, H. (2021) Production of functional oocytes requires maternally expressed PIWI genes and piRNAs in golden hamsters. bioRxiv doi: https://doi.org/10.1101/2021.01.27.428354

32 Gou, L.T., Kang, J.Y., Dai, P., Wang, X., Li, F., Zhao, S., Zhang, M., Hua, M.M., Lu, Y., Zhu, Y. et al. (2017) Ubiquitination-Deficient Mutations in Human Piwi Cause Male Infertility by Impairing Histone-to-Protamine Exchange during Spermiogenesis. Cell, 169, 1090–1104 e1013.

33 Ivanova, I., Much, C., Di Giacomo, M., Azzi, C., Morgan, M., Moreira, P.N., Monahan, J., Carrieri, C., Enright, A.J. and O’Carroll, D. (2017) The RNA m(6)A Reader YTHDF2 Is Essential for the Post-transcriptional Regulation of the Maternal Transcriptome and Oocyte Competence. Mol Cell, 67, 1059–1067 e1054.

34 Zoch, A., Auchynnikava, T., Berrens, R.V., Kabayama, Y., Schopp, T., Heep, M., Vasiliauskaite, L., Perez-Rico, Y.A., Cook, A.G., Shkumatava, A. et al. (2020) SPOCD1 is an essential executor of piRNA-directed de novo DNA methylation. Nature, 584, 635–639.

35 Raz, E. (2000) The function and regulation of vasa-like genes in germ-cell development. Genome Biol, 1, REVIEWS1017.

36 Yuan, L., Liu, J.G., Zhao, J., Brundell, E., Daneholt, B. and Hoog, C. (2000) The murine SCP3 gene is required for synaptonemal complex assembly, chromosome synapsis, and male fertility. Mol Cell, 5, 73–83.

37 Costoya, J.A., Hobbs, R.M., Barna, M., Cattoretti, G., Manova, K., Sukhwani, M., Orwig, K.E., Wolgemuth, D.J. and Pandolfi, P.P. (2004) Essential role of Plzf in maintenance of spermatogonial stem cells. Nat Genet, 36, 653–659.

38 De Fazio, S., Bartonicek, N., Di Giacomo, M., Abreu-Goodger, C., Sankar, A., Funaya, C., Antony, C., Moreira, P.N., Enright, A.J. and O’Carroll, D. (2011) The endonuclease activity of Mili fuels piRNA amplification that silences LINE1 elements. Nature, 480, 259–263.

39 Vasiliauskaite, L., Berrens, R.V., Ivanova, I., Carrieri, C., Reik, W., Enright, A.J. and O’Carroll, D. (2018) Defective germline reprogramming rewires the spermatogonial transcriptome. Nature Structural & Molecular Biology, 25, 394-+.

40 Ballow, D., Meistrich, M.L., Matzuk, M. and Rajkovic, A. (2006) Sohlh1 is essential for spermatogonial differentiation. Developmental biology, 294, 161–167.

41 Gutti, R.K., Tsai-Morris, C.H. and Dufau, M.L. (2008) Gonadotropin-regulated testicular helicase (DDX25), an essential regulator of spermatogenesis, prevents testicular germ cell apoptosis. J Biol Chem, 283, 17055–17064.

42 Li, H., Liang, Z., Yang, J., Wang, D., Wang, H., Zhu, M., Geng, B. and Xu, E.Y. (2019) DAZL is a master translational regulator of murine spermatogenesis. Natl Sci Rev, 6, 455–468.

43 Di Giacomo, M., Comazzetto, S., Sampath, S.C. and O’Carroll, D. (2014) G9a co-suppresses LINE1 elements in spermatogonia. Epigenetics & Chromatin, 7.

44 Di Giacomo, M., Comazzetto, S., Saini, H., De Fazio, S., Carrieri, C., Morgan, M., Vasiliauskaite, L., Benes, V., Enright, A.J. and O’Carroll, D. (2013) Multiple Epigenetic Mechanisms and the piRNA Pathway Enforce LINE1 Silencing during Adult Spermatogenesis. Molecular Cell, 50, 601–608.

45 Gurumurthy, C.B., Sato, M., Nakamura, A., Inui, M., Kawano, N., Islam, M.A., Ogiwara, S., Takabayashi, S., Matsuyama, M., Nakagawa, S. et al. (2019) Creation of CRISPR-based germline-genome-engineered mice without ex vivo handling of zygotes by i-GONAD. Nat Protoc, 14, 2452–2482.

46 Shatzkes, K., Teferedegne, B. and Murata, H. (2014) A simple, inexpensive method for preparing cell lysates suitable for downstream reverse transcription quantitative PCR. Sci Rep, 4, 4659.

47 Dobin, A., Davis, C.A., Schlesinger, F., Drenkow, J., Zaleski, C., Jha, S., Batut, P., Chaisson, M. and Gingeras, T.R. (2013) STAR: ultrafast universal RNA-seq aligner. Bioinformatics, 29, 15–21.

48 Teissandier, A., Servant, N., Barillot, E. and Bourc’his, D. (2019) Tools and best practices for retrotransposon analysis using high-throughput sequencing data. Mob DNA, 10, 52.

49 Liao, Y., Smyth, G.K. and Shi, W. (2014) featureCounts: an efficient general purpose program for assigning sequence reads to genomic features. Bioinformatics, 30, 923–930.

50 Love, M.I., Huber, W. and Anders, S. (2014) Moderated estimation of fold change and dispersion for RNA-seq data with DESeq2. Genome Biol, 15, 550.

51 Graf, A., Krebs, S., Zakhartchenko, V., Schwalb, B., Blum, H. and Wolf, E. (2014) Fine mapping of genome activation in bovine embryos by RNA sequencing. Proc Natl Acad Sci U S A, 111, 4139–4144.

52 Gao, Y., Li, S., Lai, Z., Zhou, Z., Wu, F., Huang, Y., Lan, X., Lei, C., Chen, H. and Dang, R. (2019) Analysis of Long Non-Coding RNA and mRNA Expression Profiling in Immature and Mature Bovine (Bos taurus) Testes. Front Genet, 10, 646.

53 Hendrickson, P.G., Dorais, J.A., Grow, E.J., Whiddon, J.L., Lim, J.W., Wike, C.L., Weaver, B.D., Pflueger, C., Emery, B.R., Wilcox, A.L. et al. (2017) Conserved roles of mouse DUX and human DUX4 in activating cleavage-stage genes and MERVL/HERVL retrotransposons. Nat Genet, 49, 925–934.

54 Jegou, B., Sankararaman, S., Rolland, A.D., Reich, D. and Chalmel, F. (2017) Meiotic Genes Are Enriched in Regions of Reduced Archaic Ancestry. Mol Biol Evol, 34, 1974–1980.

55 Horvat, F., Fulka, H., Jankele, R., Malik, R., Jun, M., Solcova, K., Sedlacek, R., Vlahovicek, K., Schultz, R.M. and Svoboda, P. (2018) Role of Cnot6l in maternal mRNA turnover. Life Sci Alliance, 1, e201800084.

56 Yue, F., Cheng, Y., Breschi, A., Vierstra, J., Wu, W., Ryba, T., Sandstrom, R., Ma, Z., Davis, C., Pope, B.D. et al. (2014) A comparative encyclopedia of DNA elements in the mouse genome. Nature, 515, 355–364.

57 Ganesh, S., Horvat, F., Drutovic, D., Efenberkova, M., Pinkas, D., Jindrova, A., Pasulka, J., Iyyappan, R., Malik, R., Susor, A. et al. (2020) The most abundant maternal lncRNA Sirena1 acts post-transcriptionally and impacts mitochondrial distribution. Nucleic Acids Res, 48, 3211–3227.

58 Yu, Y., Zhao, C., Su, Z., Wang, C., Fuscoe, J.C., Tong, W. and Shi, L. (2014) Comprehensive RNA-Seq transcriptomic profiling across 11 organs, 4 ages, and 2 sexes of Fischer 344 rats. Sci Data, 1, 140013.

59 Kent, W.J., Zweig, A.S., Barber, G., Hinrichs, A.S. and Karolchik, D. (2010) BigWig and BigBed: enabling browsing of large distributed datasets. Bioinformatics, 26, 2204–2207.

60 Gan, H., Lin, X., Zhang, Z., Zhang, W., Liao, S., Wang, L. and Han, C. (2011) piRNA profiling during specific stages of mouse spermatogenesis. RNA, 17, 1191–1203.

61 Wagih, O. (2017) ggseqlogo: a versatile R package for drawing sequence logos. Bioinformatics, 33, 3645–3647.

62 Smit, A.F.A., Hubley, R. and Green, P. (2013–2015).

63 Krueger, F. and Andrews, S.R. (2011) Bismark: a flexible aligner and methylation caller for Bisulfite-Seq applications. Bioinformatics, 27, 1571–1572.

64 Ono, M., Toh, H., Miyata, T. and Awaya, T. (1985) Nucleotide sequence of the Syrian hamster intracisternal A-particle gene: close evolutionary relationship of type A particle gene to types B and D oncovirus genes. Journal of virology, 55, 387–394.

65 Kuff, E.L. and Lueders, K.K. (1988) The intracisternal A-particle gene family: structure and functional aspects. Adv Cancer Res, 51, 183–276.

66 Kozak, C.A., Hartley, J.W. and Morse, H.C., 3rd. (1984) Laboratory and wild-derived mice with multiple loci for production of xenotropic murine leukemia virus. Journal of virology, 51, 77–80.

67 Costas, J. (2003) Molecular characterization of the recent intragenomic spread of the murine endogenous retrovirus MuERV-L. Journal of molecular evolution, 56, 181–186.

68 Kigami, D., Minami, N., Takayama, H. and Imai, H. (2003) MuERV-L is one of the earliest transcribed genes in mouse one-cell embryos. Biol Reprod, 68, 651–654.

69 Macfarlan, T.S., Gifford, W.D., Agarwal, S., Driscoll, S., Lettieri, K., Wang, J., Andrews, S.E., Franco, L., Rosenfeld, M.G., Ren, B. et al. (2011) Endogenous retroviruses and neighboring genes are coordinately repressed by LSD1/KDM1A. Genes Dev, 25, 594–607.

70 Davis, M.P., Carrieri, C., Saini, H.K., van Dongen, S., Leonardi, T., Bussotti, G., Monahan, J.M., Auchynnikava, T., Bitetti, A., Rappsilber, J. et al. (2017) Transposon-driven transcription is a conserved feature of vertebrate spermatogenesis and transcript evolution. EMBO Rep, 18, 1231–1247.

71 Brouha, B., Schustak, J., Badge, R.M., Lutz-Prigge, S., Farley, A.H., Moran, J.V. and Kazazian, H.H. Jr. (2003) Hot L1s account for the bulk of retrotransposition in the human population. Proc Natl Acad Sci U S A, 100, 5280–5285.

