## Supplementary figures and images for "Golden hamster piRNAs are necessary for early embryonic development and establishment of spermatogonia"

### Data S1ERVK LTR retrotransposon nucleotide exchange rates

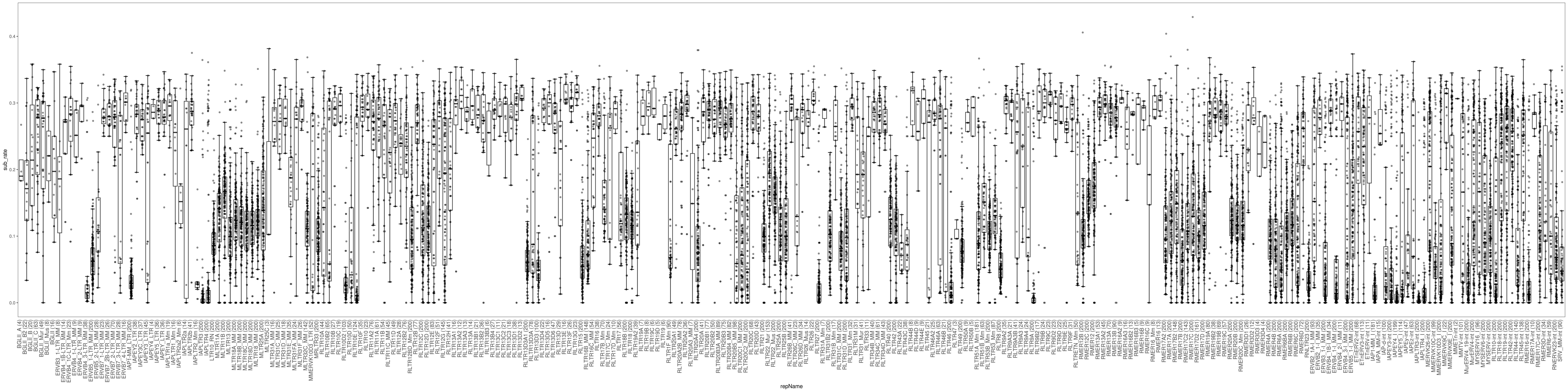

### Data S2Other LTR retrotransposon nucleotide exchange rates

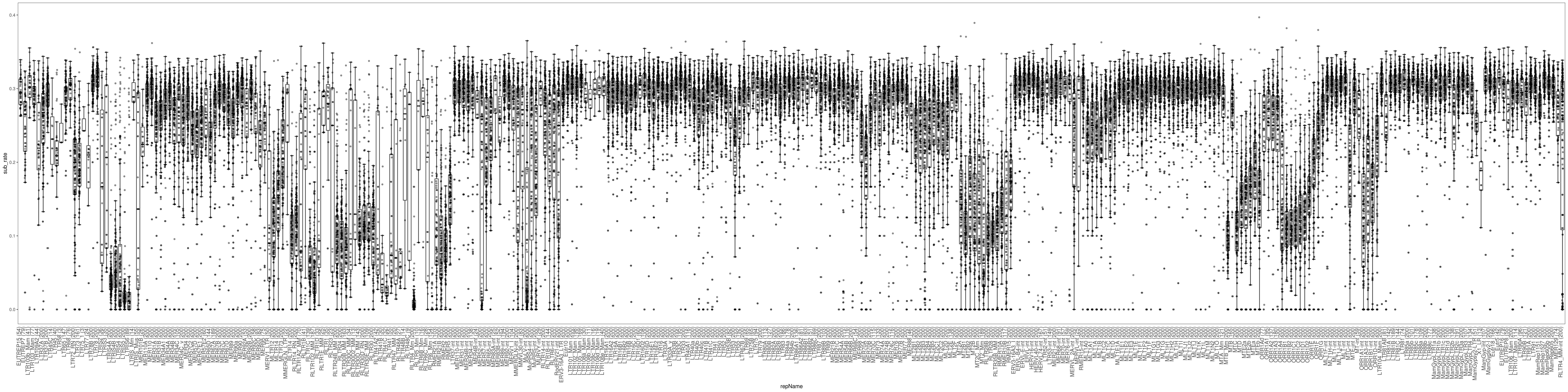

### Data S3MaLR LTR retrotransposon nucleotide exchange rates

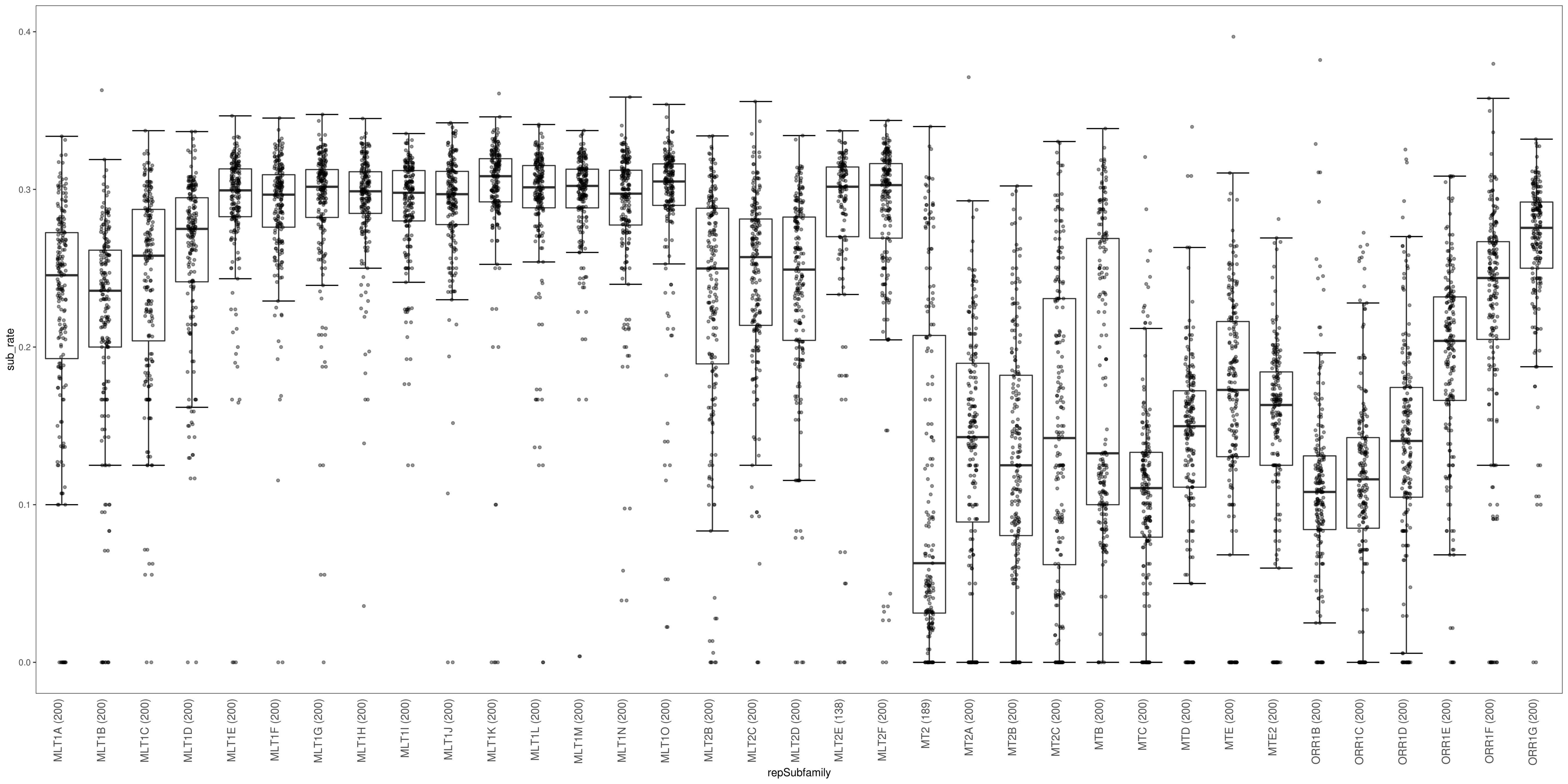
